# Theta oscillations coincide with sustained hyperpolarization in CA3 pyramidal cells, underlying decreased firing

**DOI:** 10.1101/652461

**Authors:** Meryl Malezieux, Ashley L. Kees, Christophe Mulle

**Author notes:** These authors contributed equally (Meryl Malezieux, Ashley L. Kees).

## Abstract

Brain-state fluctuations modulate membrane potential dynamics of neurons, influencing the functional repertoire of the network. Pyramidal cells (PCs) in hippocampal CA3 are necessary for rapid memory encoding, preferentially occurring during exploratory behavior in the high-arousal theta state. However, the relationship between the membrane potential dynamics of CA3 PCs and theta has not been explored. Here, we characterize the changes in the membrane potential of PCs in relation to theta using electrophysiological recordings in awake mice. During theta, most PCs behave in a stereotypical manner, consistently hyperpolarizing time-locked to the duration of theta. Additionally, PCs display lower membrane potential variance and reduced firing rate. In contrast, during large irregular activity, a low-arousal state, PCs show heterogeneous changes in membrane potential. This suggests coordinated hyperpolarization of PCs during theta, possibly caused by increased inhibition. This could lead to higher signal-to-noise ratio in the small population of PCs active during theta as observed in ensemble recordings.

## Introduction

Wakefulness is comprised of distinct brain states, correlated with different behaviors and characterized by specific oscillatory patterns in the local field potential (LFP). The ability of brain circuits to perform the different computations that give rise to different behaviors is thought to rely upon state-dependent modulations in single-cell properties, such as resting membrane potential, spike threshold, and synaptic efficacy (Marder et al., 2014), and network-level properties such as upstream inputs (Buzsáki, 2002). By providing a window into both the single-cell properties and the integration of synaptic inputs into firing output, single-cell membrane potential recordings *in vivo* in awake animals are essential to our understanding of the relationship between neuronal computation and brain states.

The hippocampus integrates multimodal information and plays a critical role in the encoding, consolidation and retrieval of episodic memories (Buzsáki and Moser, 2013; Eichenbaum, 2016). A large number of studies using extracellular recordings in freely behaving animals have demonstrated that neural ensembles in the hippocampus support representations of space and other task-relevant stimuli (Buzsáki and Moser, 2013). Different brain states are associated with distinct sensory integration properties as well as specific memory functions. Exploratory behaviors are associated with robust theta (4–12 Hz) and gamma (30–80 Hz) oscillations in the LFP. Since their discovery in the rabbit hippocampus, theta rhythms have been extensively studied and found in many species including rats, mice, monkeys and humans (Ekstrom et al., 2005; Green and Arduini, 1954; Jutras et al., 2013; Vanderwolf, 1969). The percentage of time in which theta is present in the LFP can predict the rate of acquisition and the quality of a memory, suggesting an important role for theta in the formation of memories (Berry and Thompson, 1978; Gupta et al., 2012; Landfield et al., 1972; Mizuseki et al., 2009). Additionally, theta correlates with behaviors associated with active sampling of the environment, like exploring, sniffing and whisking, and therefore appears well suited for the coordination of multimodal sensory integration coming from the entorhinal cortex (Komisaruk, 1970; Macrides et al., 1982). During non-locomotor behaviors, such as immobility, eating, drinking, and grooming, as well as during slow wave sleep, large irregular activity (LIA) can be detected in the hippocampal LFP. The LIA pattern is marked by high-amplitude broadband fluctuations rather than oscillations within a specific frequency band and is often characterized by the presence of sharp-waves (40-100 ms duration) that co-occur with high-frequency ripple oscillations (80-250 Hz) (SWRs) in the CA1 LFP (Buzsáki, 1986; O’Keefe, 1976; Vanderwolf, 1969). LIA is present during periods of low arousal and is thought to represent a state in which memory consolidation and retrieval of episodic memory is promoted (Buzsáki, 2015; Carr et al., 2011).

Membrane potential recordings from single CA1 pyramidal cells (PCs) in awake behaving animals has provided valuable insight into the relationship between network and single-cell activity and its role in behavior and cognition. The combination of intracellular and extracellular recordings *in vivo* in awake rodents has revealed synaptic mechanisms of CA1 PCs SWRs (English et al., 2014; Gan et al., 2017; Hulse et al., 2016; Maier et al., 2011) and allowed characterization of membrane potential dynamics in relation to brain state and arousal (Hulse et al., 2017). CA1 hippocampal place cells have been recorded intracellularly in both freely moving (Epsztein et al., 2011) and head-fixed (Bittner et al., 2015; Cohen et al., 2017; Harvey et al., 2009) rodents, enabling empirical testing of the neural computations underlying spatial cognition.

While much has been learned about the relationship between hippocampal LFP oscillations and single-cell properties in CA1, its main input source, CA3, remains largely under-investigated. The few studies that have examined the intracellular activity of CA3 PCs *in vivo* have been performed in anesthetized rats (Atallah and Scanziani, 2009; Kowalski et al., 2016) and mice (Zucca et al., 2017), therefore precluding the study of awake brain states. CA3 is unique in both its anatomical and functional characteristics; CA3 PCs display extensive recurrent connections, and receive strong mossy fiber input from dentate granule cells (Rebola et al., 2017). Computational theories have long proposed that CA3 rapidly stores memories through synaptic plasticity in the autoassociative network formed by its recurrent connections (Kesner and Rolls, 2015), and CA3 is indeed necessary for one-trial memory encoding (Nakazawa et al., 2003). Moreover, theory and experimental evidence support that efficient episodic memory representations require a sparsely distributed neural code, in which each memory is coded by the activity of a small proportion of hippocampal neurons (Leutgeb et al., 2007; Marr, 1971; Wixted et al., 2014). To support their various functions, CA3 PCs must dynamically adapt their intracellular properties to the ongoing behavioural and brain state of the animal. To explore the modulation of intracellular dynamics of CA3 PCs in relation to changes in brain states, we performed whole-cell patch-clamp and LFP recordings in awake head-fixed mice in combination with measurements of pupil diameter and running activity.

We find that theta is characterized by the hyperpolarization of most CA3 PCs, accompanied by a decrease in their firing rate. Additionally, the amplitude of subthreshold membrane potential fluctuations decreases during theta. In contrast, we observed heterogeneous changes in membrane potential dynamics during LIA. Altogether, we show that several single-cell and network-level modulations are taking place during theta, impacting firing output of CA3 pyramidal cells, and we propose mechanisms underlying these changes.

## Results

### Hippocampal CA3 LFP characterization and brain state classification

To investigate the intracellular dynamics of CA3 PCs *in vivo* across brain states, we performed simultaneous whole-cell and LFP recordings in awake head-fixed mice free to run on a wheel (Figure 1A). An LFP electrode was first positioned in the *stratum pyramidale* of dorsal CA3 before lowering the patch-clamp electrode for whole-cell recordings in the vicinity of the LFP electrode. The tips of the LFP and patch electrodes were not further than 200 µm apart, as confirmed by posthoc histology. Pyramidal cell identity was determined by the voltage response to current steps as well as by the presence of spontaneous or evoked burst firing (Figure 1B and 1C). To monitor the level of arousal, we measured both pupil diameter and locomotor activity on the wheel (Figures 1A and 1C). Figure 1C shows an example of a current-clamp recording from a CA3 PC aligned with the corresponding LFP recording, locomotor activity and pupil diameter. We recorded a total of 33 CA3 PCs in 21 animals. In conjunction with the electrophysiological criteria for cell identity described above, all recordings were confirmed to have patch pipette tracks in CA3. Moreover, for 14 recordings, we were able to recover the cell body location with biocytin, further confirming their identity (Figure 1D). Recordings were discarded when the location was unable to be confirmed in CA3. The basic properties of each CA3 PC recorded are shown in Tables S1 and S2.

**Figure 1.**
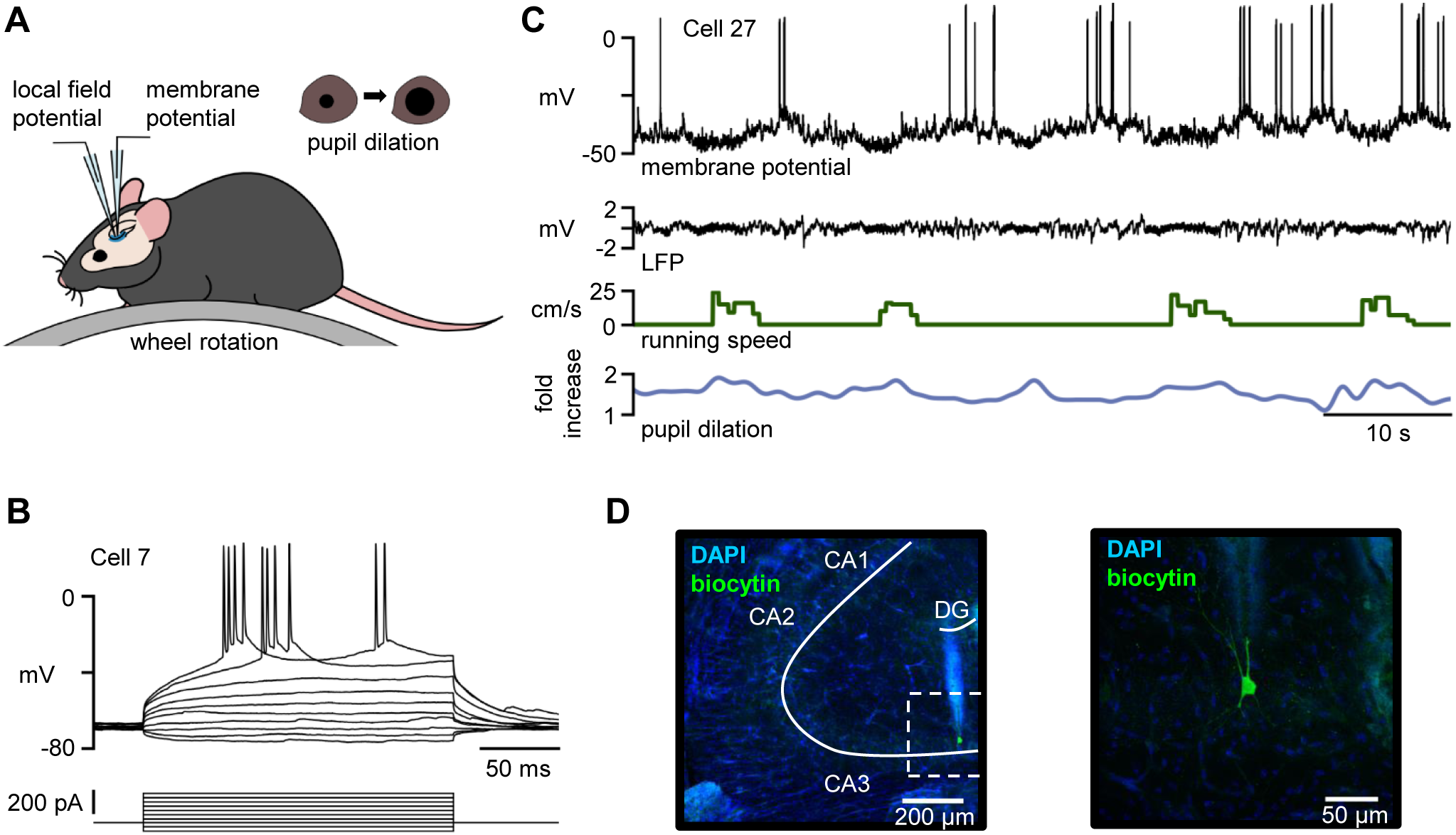
Experimental configuration and data recorded. (A) Schematic of the head-fixed electrophysiological recording configuration. LFP and whole-cell recordings were performed in CA3. Wheel rotation and pupil dilation were monitored simultaneously. (B) Example trace of the sub- and suprathreshold responses of a CA3 PC to injected current pulses (10 pulses, from -80 pA to 280 pA). (C) Example trace of a simultaneous current-clamp recording from a CA3 PC showing spontaneous action potentials (membrane potential, black), aligned with the nearby LFP (LFP, black) the corresponding locomotor velocity (running speed, green) and pupil diameter changes (pupil dilation, blue) of the animal. Cell id numbers correspond to Tables S1 and S2. (D) Confocal fluorescent image of a 100-µm-thick coronal section of dorsal hippocampus (left; 20x) with a single CA3 PC (right; 40x) filled with biocytin during whole-cell recording and visualized by post hoc labeling with Alexa Fluor 488. Note the presence of the pipette track above the filled CA3 PC.

Brain state classification was achieved offline using the spectral properties of the LFP to find epochs of theta and LIA (Figures 2A and 2B), using methods similar to those previously described (Hulse et al., 2017). An event was defined as a single epoch of theta or LIA, with a defined onset and offset. Theta was detected using the ratio of power in the theta (6-9 Hz) range to power in the delta (0.5-3 Hz) range (Figure 2B). From a total of 695 theta events detected from all recordings, 52% occurred in conjunction with detected locomotor activity (run theta); the remaining 48% were categorized as rest theta. We cannot exclude that rest theta was correlated with finer exploratory movements of the mouse such as whisking and sniffing which we did not monitor. LIA was detected as peaks in the average broadband power (0.3-80 Hz) after z-scoring within frequency bands; since some peaks in the broadband power were due to an increase in only the theta band, events that overlapped with detected theta events were not included (Figure 2B). Theta and LIA events were dispersed throughout the session with no apparent pattern (Figure 2C), and all three types of events occurred with equal frequency (Figure 2D). Out of the three event types, run theta occurred for the longest duration (Figure 2E). Over a total recording time of 223 minutes, 12% of the time was classified as run theta, 7% as rest theta, and 11% as LIA (Figure 2F). The pupil diameter of the mice was smaller during LIA as compared to both types of theta events (Figures 2G and S1). This is consistent with previous reports (Hulse et al., 2017), and provides an independent confirmation of the criteria used in the detection of brain states. Interestingly, the theta-delta ratio was significantly higher during run theta than rest theta (Figure 2H). For run theta events, increases in the theta-delta ratio often preceded the start of the detected run (Figure 2I), which has also been noted in previous studies (Fuhrmann et al., 2015; Green and Arduini, 1954; Vanderwolf, 1969). Thus, our stringent criteria for theta and LIA events allowed us to proceed to characterize the intracellular dynamics of CA3 PCs in relation to the brain state.

**Figure 2.**
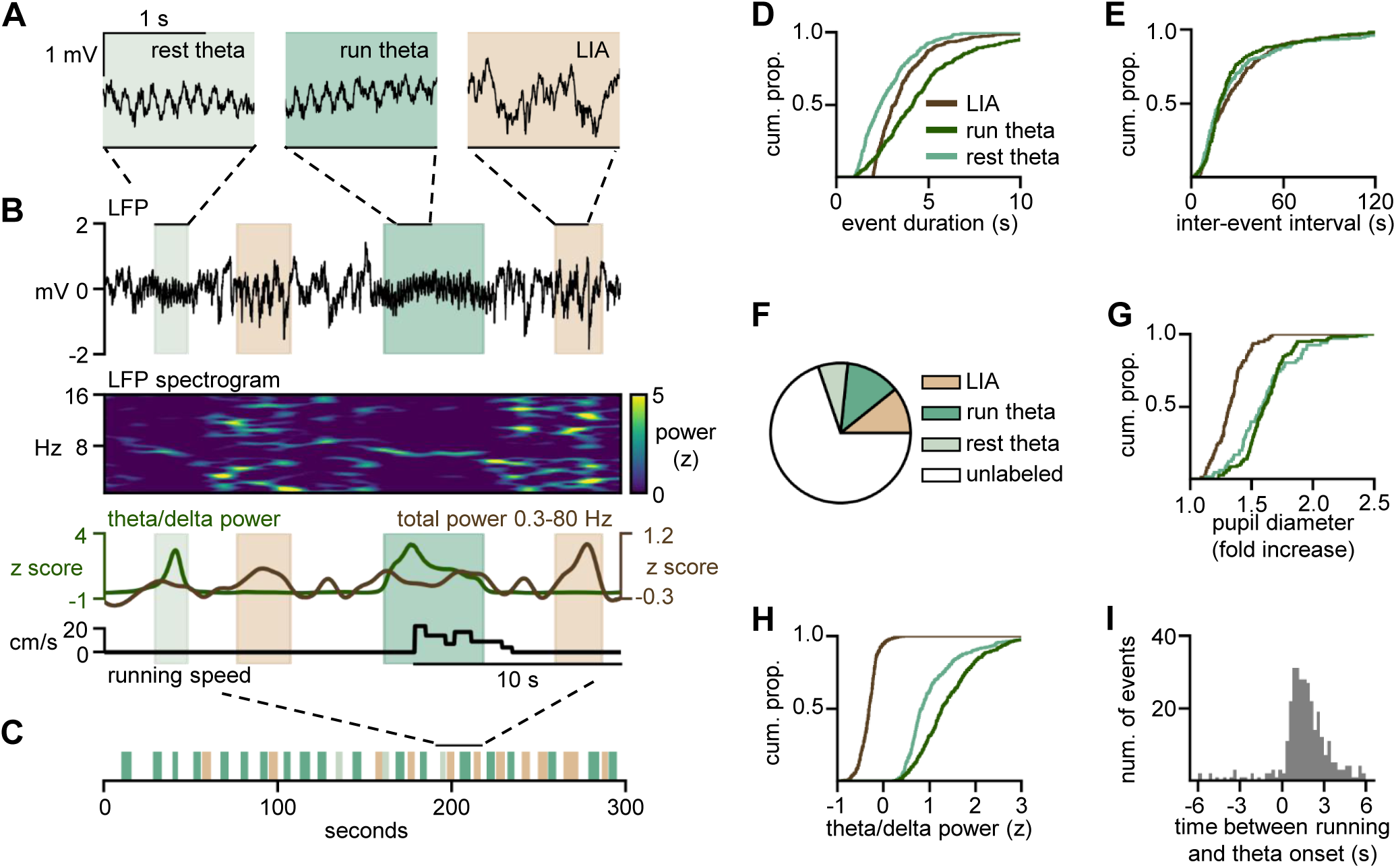
Brain state classification and characterization. (A) Example traces of rest theta (light green), run theta (dark green) and LIA (brown). Similar color scheme is used throughout. (B) Top: LFP recording from CA3 with detected LIA, run theta and rest theta. Middle: Corresponding spectrogram showing z-scored LFP power between 0 and 16 Hz. Note the high broadband power present during LIA and narrow frequency band during theta events. Bottom: Theta/delta power ratio superimposed with total 0.3-80 Hz power, used in state detection. Velocity of the animal is shown below. (C) Representation of the brain states LIA, run theta and rest theta over a full recording session. (D) Cumulative distribution of the duration of the different states. Run theta events lasted longer than rest theta and LIA events (4.2 ± 2.0 s [n = 362 events] for run theta, 2.4 ± 1.1 s [n = 333 events] for rest theta, 3.2 ± 1.2 s [n = 397 events] for LIA, p < 0.001 for both pairs). LIA events lasted longer than rest theta (p < 0.001). All data are expressed in median ± average absolute deviation from the median unless otherwise stated. (E) Cumulative distribution of the inter-event interval of the different states. All event types occurred at a similar frequency (Kruskal-Wallace; p = 0.2) (19.3 ± 21.5 s [n = 362 events] for run theta, 19.1 ± 20.0 s [n = 333 events] for rest theta, 22.5 ± 20.0 s [n = 397 events] for LIA). (F) Total percent of time detected for each state over all the recording sessions (n = 33 recordings). Over 223.7 minutes of recording there were a total of 28.2 minutes of run theta (12%), 15.3 minutes of rest theta (7%), and 24.0 minutes of LIA (11%). The remaining 156.2 minutes (70%) were left unlabeled. (G) Pupil diameter (expressed as fold increase from the minimum) during each brain state (n = 12 recordings). The pupil dilated the same amount for run and rest theta (p = 0.682), but was more dilated during both types of theta events than during LIA (1.6 ± 0.2 fold [n = 117 events] for run theta, 1.6 ± 0.2 fold [n = 66 events] for rest theta, 1.3 ± 0.1 fold [n = 77 events] for LIA, p < 0.001 for both pairs). (H) Theta/delta power ratio for each brain state. The z-scored theta/delta power was larger during theta than during LIA (1.3 ± 0.5 [n = 362 events] for run theta, 0.9 ± 0.5 [n = 333 events] for rest theta, -0.3 ± 0.2 [n = 397 events] for LIA, p < 0.001 for both pairs). This ratio was also larger during run theta compared to rest theta (p < 0.001). (I) Delay between the start of theta and start of running. Running started significantly after the start of theta (1.5 ± 6.8 s, p < 0.001 [n = 360 events]).

### Theta is characterized by hyperpolarization of CA3 PCs

We explored the membrane potential of CA3 PCs in relation to theta (Figure 3), a high-arousal brain state during which active sampling of the environment and integration of stimuli occur. We performed our analyses on smoothed and downsampled membrane potential traces after spike removal. We observed that the membrane potential hyperpolarized at the onset of a large proportion of theta events (Figure 3A and S2). To quantify this hyperpolarization on an event-by-event basis, we compared windows before and after theta onset that aligned with the maximal change in the average trace (Figure S2B) (-2.5 to -0.5 seconds and 0.5 to 2.5 seconds). The magnitude of the change was defined as the maximal difference between the potentials in the two windows, and the significance was determined using a Welch’s t-test. Out of 668 theta events, 52% were associated with a significant hyperpolarization, 19% with depolarization and 29% showed no significant modulation (Figure 3B). These proportions were not different between run theta and rest theta, suggesting that the relationship between theta and membrane potential is more related to the underlying brain state than the overt behavior (Figure S2C). Over the population of cells, there was a significantly higher proportion of hyperpolarizing events and significantly lower proportion of depolarizing and nonsignificant events compared with randomly selected times. Since this analysis could be biased by a few cells with many theta events, we confirmed the relationship between theta and hyperpolarization by performing a bootstrap analysis on each cell individually. For each cell, we created shuffled data sets by advancing the time of the theta events with respect to the membrane potential trace; cells that showed a larger positive or negative average change than was observed in the 95% of the shuffled data were considered significantly depolarizing or hyperpolarizing, respectively. By this measure, 17 cells were significantly hyperpolarizing during theta, 1 was significantly depolarizing, and 15 showed no difference with the shuffled data set, perhaps due to a variable change in membrane potential at the onset of theta events (Figure 3C). We did not find any significant difference in access resistance, input resistance, or capacitance between cells that consistently hyperpolarized during theta and those that did not (Table S1). We observed a tight correlation between the duration of hyperpolarization and the duration of theta events. Indeed the membrane potential was hyperpolarized for the whole duration of the theta event and returned to baseline close to the offset of the theta event (Figure 3D and 3E). Depolarizations, on the other hand, started together with the theta event but seemed to persist beyond the end of the theta event (Figure 3D and 3E).

**Figure 3.**
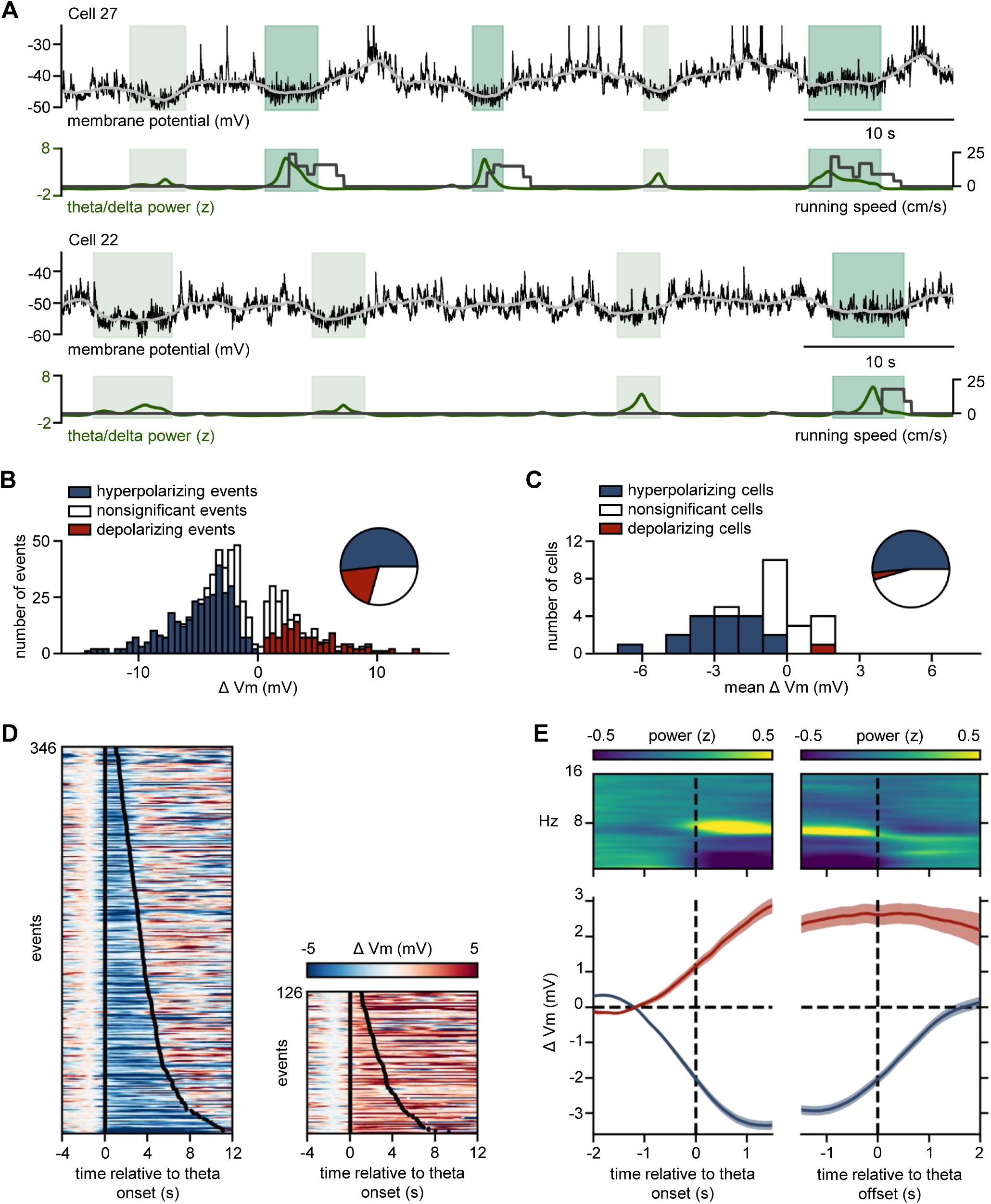
CA3 PCs hyperpolarize during theta. (A) Example whole-cell recordings from two different CA3 PCs (top), with simultaneous theta/delta power ratio of the nearby LFP and the running speed of the mouse (bottom). Grey trace superimposed on the raw Vm is the smoothed Vm after spike removal, and green shading represents theta events. Spikes are truncated in the raw membrane potential traces. (B) Distribution of the magnitude of change in Vm at the onset of theta (n = 668 events, bin size = 0.5 mV). Events are colored according to whether there was a hyperpolarization (blue), depolarization (red) or no significant change (white). Similar color scheme is used throughout the figures. Inset: pie chart of the different event types (52% hyperpolarizing [n = 346 events], 19% depolarizing [n = 126 events], 29% no change [n = 196 events]). Compared to a shuffled data set, there were significantly more hyperpolarizing events and fewer depolarizing and nonsignificant events (p < 0.001 for all categories) than chance level. (C) Distribution of cells’ average Vm change during theta (bin size = 1 mV). Inset: pie chart of the different cell types (17 cells were hyperpolarizing, 1 cell was depolarizing and 15 cells showed no significant modulation towards one specific direction). (D) Color plots showing the Vm of CA3 PCs triggered by transition to theta normalized to 2.5 to0.5 s before start of theta. Blue represents Vm hyperpolarization, red Vm depolarization. Events are separated whether they are associated with a significant hyperpolarization (left, n = 346 events), or depolarization (right, n = 126 events). Events with no change in Vm are not shown (n = 196 events). Events are ordered by their length, and the black lines show the start and stop of events. (E) Average spectrograms and membrane potentials triggered by theta onset (left) and offset (right). Top: Average spectrogram. Bottom: Average membrane potential traces (after spike removal and smoothing) for events with hyperpolarization (blue) and depolarization (red). Shaded portions represent ± SEM.

We then asked if the observed modulation of membrane potential of CA3 PCs during theta have an impact on their firing properties (Figure 4A). Indeed, significant changes in membrane potential had a corresponding change in average firing rate, both in individual events and in cell averages (Figure 4B-E). No difference was observed in the properties of complex spikes (CS) or the proportion of spikes occurring within them (Figure S3). Taken together, these data demonstrate a robust hyperpolarization and correlated decrease in firing rate during theta in CA3 pyramidal cells.

**Figure 4.**
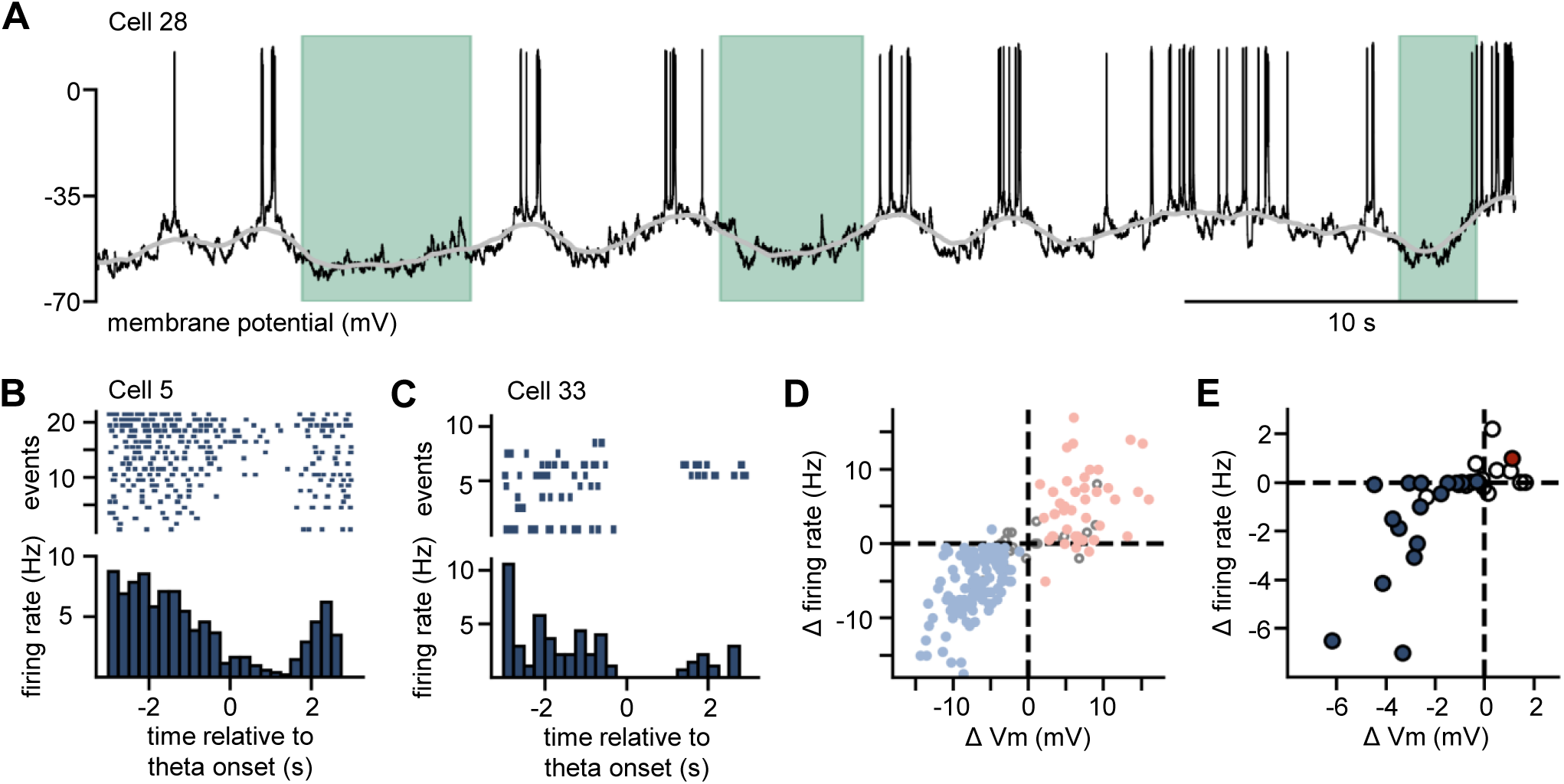
Firing rate of CA3 PCs decreases during theta. (A) Example trace of a theta-hyperpolarizing cell. Shaded areas are theta events. (B) and (C) Firing rate around the start of theta for two example theta hyperpolarizing cells. Top: raster plot where a single tick represents a single spike and each row is a single transition to theta triggered at time 0. Bottom: PSTH representing the average firing rate of the same cell for all theta transitions triggered at time 0. (D) Change in firing rate versus change in Vm during theta events (n = 194 events). Filled dots represent significant Vm modulation during theta, color represents the direction of modulation. (E) Mean change in firing rate versus mean change in Vm during theta for each cell (n = 33 cells). Filled dots represent significant Vm modulation during theta, color represents the direction of modulation.

A straightforward explanation of the above results is that hyperpolarization brings the cell further from a stable firing threshold, leading to a decrease in firing rate. However, we found that the data were not fully consistent with this hypothesis. Specifically, we found that in individual cells, the thresholds of spikes (see methods) varied by several mV over short time scales (Figure S4A). This was correlated with the baseline membrane potential before the spike, suggesting that the spike threshold adapts to slow changes in the membrane potential and counteracts the tendency of the cell to fire (Azouz and Gray, 2003; Epsztein et al., 2011). The distance between baseline membrane potential and threshold decreases at depolarized membrane potentials, suggesting that slow changes in membrane potential have some ability to bring the cell closer to spike threshold (Figure S4B). This distance between membrane potential and threshold was not different during theta as compared with unlabeled states (Figure S4C), suggesting that the hyperpolarizations were insufficient to bring the cell further from spike threshold. Next, we looked at the pre-spike depolarization (PSD; see methods) (Poulet and Petersen, 2008), a measure to assess the amount of coincident input necessary to elicit a spike. We did not find any theta-dependent changes (Figure S4D), suggesting that changes in firing rate were not due to changes in excitability. These results suggest that absolute threshold varies dynamically for a given cell, and that the distance between ongoing membrane potential and absolute threshold stays similar during theta.

### LIA is associated with heterogeneous modulation of CA3 PCs

We similarly analyzed the intracellular properties of CA3 PCs during LIA (Figure 5A, Figure S5), a low-arousal brain state. Out of 378 LIA events, 45% were associated with a depolarization, 18% with a hyperpolarization, and 37% with no change, which represents more depolarizations and fewer hyperpolarizations than would be expected by chance (Figure 5B). However, when comparing each cell’s modulations with those from shifted time series, only 8 cells consistently depolarized, and 24 cells showed no consistent modulation (in one cell, no LIA events were recorded) (Figure 5C). Although LIA events are associated with a high proportion of depolarizations, it seems that those depolarizations happen throughout the cell population and not specifically in a subset of cells. Therefore, many cells have a variety of responses to different LIA events, in line with recent research showing different ensembles are active during different SWRs (Ramirez-Villegas et al., 2015; Taxidis et al., 2015). The depolarizing events and depolarizing cells were also associated with increased firing rate (Figure 5D-G). Similarly to theta, relative threshold (Figure S6A-C) and PSD (Figure S6D) was not modulated during LIA. Thus, LIA-depolarizing cells do not increase their firing rate due to a change in excitability, but most probably due to an increase in the frequency or amplitude of synaptic inputs.

**Figure 5.**
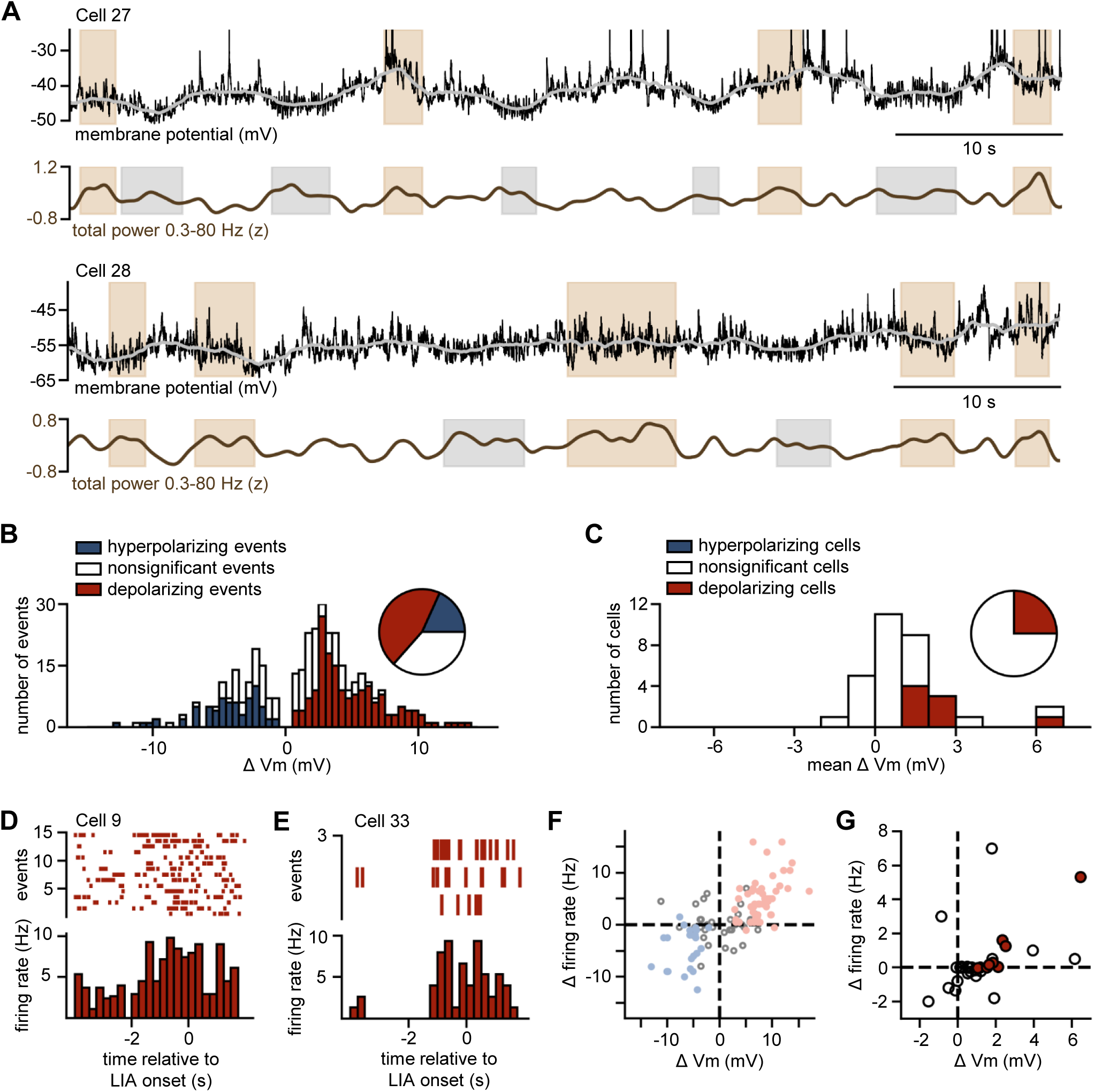
CA3 pyramidal cells have heterogeneous changes in membrane potential during LIA. (A) Example whole-cell recordings from two different CA3 PCs (top), with simultaneous total power of the nearby LFP (bottom). Grey trace superimposed on the raw Vm is the smoothed Vm after spike removal, and brown shading represents LIA events. Spikes are truncated in the raw membrane potential traces. (B) Distribution of the magnitude of change in Vm at the onset of LIA (n = 378 events, bin size = 0.5 mV). Events are colored according to whether there was a hyperpolarization (blue), depolarization (red) or no significant change (white). Inset: pie chart of the different event types (18% hyperpolarizing [n = 69 events], 45% depolarizing [n = 171 events], 37% no change [n = 138 events]). Compared to a shuffled data set, there were significantly more depolarizing events and fewer hyperpolarizing events (p < 0.001 for all categories) than chance level. (C) Distribution of cells’ average Vm change during LIA (bin size = 1 mV). Inset: pie chart of the different cell types (0 cells were hyperpolarizing, 8 cells were depolarizing and 24 cells showed no significant modulation towards one specific direction; 1 cell did not have any LIA events). (D) and (E) Firing rate around the start of LIA for two example depolarizing cells. Top: raster plot where a single tick represents a single spike and each row is a single transition to LIA triggered at time 0. Bottom: PSTH representing the firing rate of the same cell for all LIA transitions triggered at time 0. (F) Change in firing rate versus change in Vm during LIA events (n = 114 events). Filled dots represent significant Vm modulation during LIA, color represents the direction of modulation. (G) Mean change in firing rate versus mean change in Vm during LIA for each cell (n = 32 cells). Filled dots represent significant Vm modulation during LIA, color represents the direction of modulation.

Of the 9 LIA-depolarizing cells, 6 were also theta-hyperpolarizing, suggesting that this is a common combination of responses and that there is considerable overlap between cells that are consistently modulated by either brain state (Figure S5C, Table S1).

### Theta-associated hyperpolarization and decreased firing rate occur with changes in input resistance and membrane potential variance

To provide a mechanistic explanation for the modulation of membrane potential observed during theta, we examined the correlation between the change in membrane potential and the initial membrane potential before the state transition (Figure 6A). To exclude the possible influence of the holding current imposed on the cell membrane, we restricted this analysis to recording times when no holding current was applied. The initial membrane potential correlated with the amplitude and direction of the change in membrane potential during theta events. Specifically, larger hyperpolarizations occurred at more depolarized initial membrane potential (Figure 6B). However, both hyperpolarizations and depolarizations occurred over the range of membrane potentials recorded, so the high incidence of hyperpolarizations was not merely due to the presence of depolarized cells. During LIA, the change in membrane potential is influenced by the initial membrane potential value to a lesser extent (Figure S7A and B).

**Figure 6.**
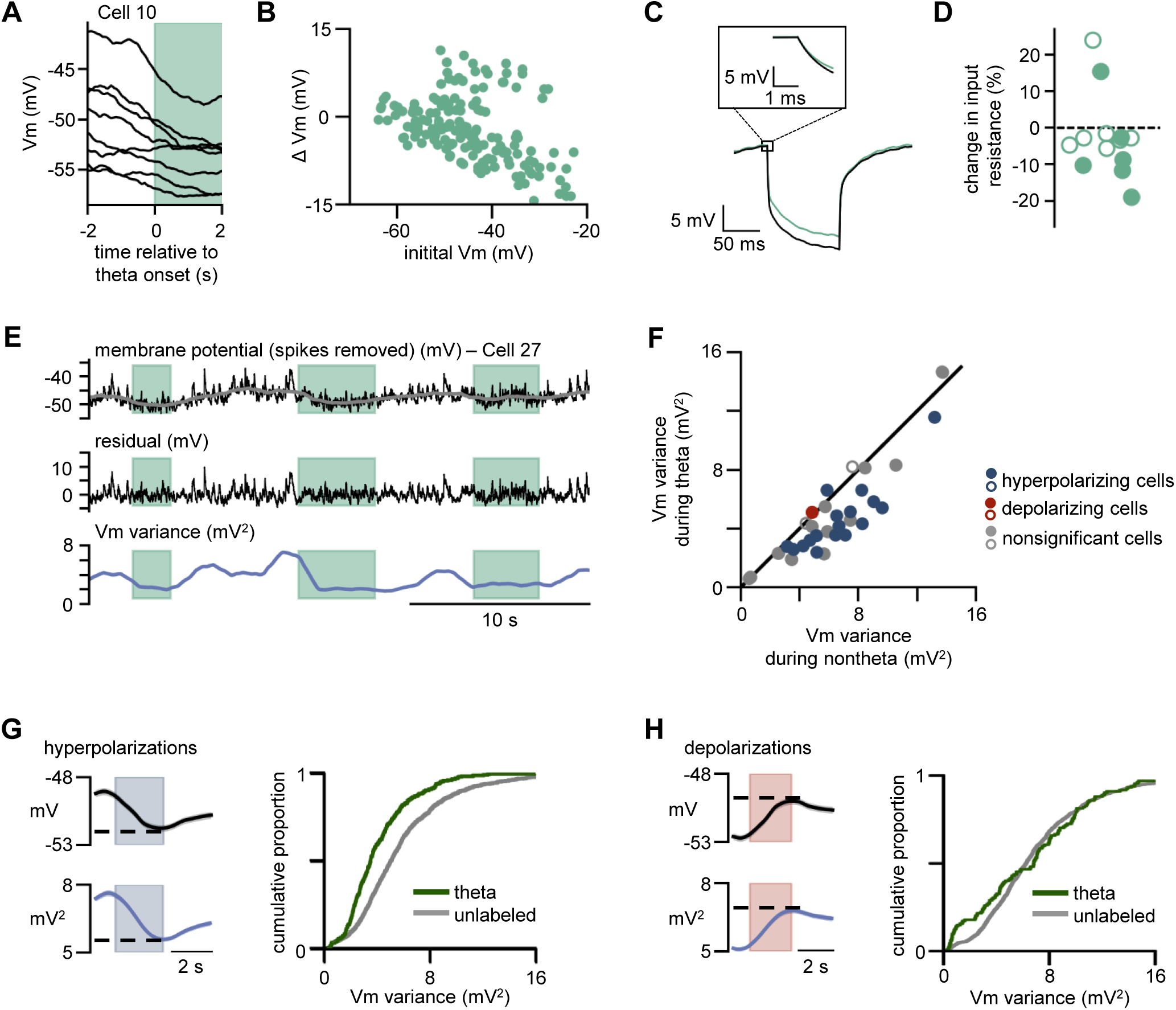
Theta-associated changes in membrane potential and firing rate occur with changes in input resistance and membrane potential variability. (A) Example traces showing Vm modulation during theta for different initial Vm values in a single cell. (B) Vm changes during theta (ΔVm) versus the initial membrane potential (initial Vm) of the cell [n = 170 events]. (C) Average voltage response to current pulses (-100 pA, 100 ms) in an example cell during theta (green) and unlabeled (black). Inset: zoom in at the onset of the current pulse. (D) Percentage change in input resistance during theta compared to unlabeled (-3.4% ± 6.9%, p = 0.491, [n = 13 cells]). Filled circles represent cells that had a significant difference in input resistance between theta and unlabeled states [n = 6 cells]. (E) Processing of Vm trace to measure its variance. Top: downsampled Vm after spike removal of a CA3 PC with shading indicating theta (green). The grey superimposed trace is after smoothing with a window of 2 seconds. The smoothed Vm is subtracted from the spike-removed Vm to obtain the residual Vm (middle). The variance of the residual Vm trace is calculated over 1-second windows (bottom). (F) Scatter plot showing Vm variance of each cell during theta versus nontheta (n = 33 cells). Over the population of cells, there was a significant decrease in variance during theta (5.8 ± 2.2 mV^2^ for nontheta, 4.2 ± 2.0 mV^2^ for theta, mean difference = -1.5 mV^2^, p < 0.001). Color represents direction of modulation during theta (blue: significantly hyperpolarizing cell: red: significantly depolarizing cell: grey: nonsignificant change), and filled dots represent a significant difference in variance between theta and nontheta. (G) Left: Average Vm (top, black) and variance (bottom, blue, n = 1625 events) of all hyperpolarizing events. Dotted line corresponds to the value taken at the end of the change in Vm for each event. Right: Cumulative proportion of variance during theta (green) and unlabeled (grey) hyperpolarizing events. Vm variance is lower during theta (3.5 ± 1.9 mV^2^ [n = 363 events] for theta, 5.1 ± 2.8 mV^2^ [n = 1177 events] for unlabeled, p < 0.001). (H) Left: Average Vm (top, black) and variance (bottom, blue, n = 1719 events) of all depolarizing events. Dotted line corresponds to the value taken at the end of the change in Vm for each event. Right: Cumulative proportion of variance during theta and unlabeled depolarizing events. Vm variance was not significantly different during depolarizations that occur during theta or unlabeled states (5.2 ± 3.8 mV^2^ [n = 139 events] for theta, 6.1 ± 3.5 mV^2^ [n = 1399 events] for unlabeled, p = 0.207).

During theta, input resistance of CA3 PCs modestly but significantly decreased in 6 out of 13 cells (Figure 6C and D). During LIA, a different set of 6 cells increased their input resistance (Figure S7C and D).

In cortical areas, high arousal states have been associated with fewer spikes and lower membrane potential variance of principal cells (Bennett et al., 2013; Lin et al., 2019; McGinley et al., 2015). To address the possibility that theta would be associated with similar changes, we measured the variance of the subthreshold membrane potential. Figure 6E shows an example of the membrane potential of a cell after spike removal (top black trace) and smoothing over windows of 2 seconds (grey trace). The residual trace was calculated by taking the difference between the spike-removed and smoothed traces; the variance was measured from this trace in sliding windows of 1 second. In comparison to all other times, the variance was significantly lower during theta and significantly higher during LIA (Figure 6F and S7F). Since the residual trace preserved all subthreshold fluctuations with a frequency higher than 0.5 Hz, this result is not due to differential filtering during the analysis of the membrane potential during theta and LIA. Given that membrane potential variance tends to correlate with the average membrane potential, we reanalyzed the data controlling for this covariate. To do this analysis, we first identified all the significant hyperpolarizing and depolarizing events (see methods) and separated them based on whether they coincided with the onset of theta or LIA. The variance was then taken at the end of each hyperpolarization or depolarization. Variance was lower during theta-associated than unlabeled-associated hyperpolarizations (Figure 6G), In contrast, the variance after a depolarization was not significantly different between those occurring at the start of theta states and those occurring outside states (Figure 6H). Taken together, these results suggest that the reduction in membrane potential variability that occurs during theta is not merely a byproduct of the change in membrane potential. In comparison, the variance observed during LIA was not different from the variance observed during depolarizations occurring outside states (Figure S7G and H). Increased synaptic input from GABAergic interneurons should normally favor hyperpolarization, with an initial increase in membrane potential variability followed by a decrease as the cell membrane approaches the reversal potential for chloride. Consistent with this hypothesis, we found a nonzero delay between hyperpolarization of the membrane potential and changes in membrane potential variability, but this was not state-dependent (Figure S8).

## Discussion

Modulations in single-cell properties, such as resting membrane potential, intrinsic excitability and synaptic activity are thought to underlie the ability of brain circuits to generate the activity patterns that correlate with distinct brain states (Marder et al., 2014). In the neocortex, recordings of single cell membrane potential dynamics across brain states and during behavior have been key in elucidating the cellular mechanisms underlying the occurrence of distinct network activity patterns (Crochet and Petersen, 2006; Polack et al., 2013; Schneider et al., 2014). In the hippocampus of awake behaving rodents, such recordings have been restricted to CA1 (English et al., 2014; Gan et al., 2017; Hulse et al., 2016; Maier et al., 2011). Here we provide for the first time data showing the intracellular dynamics of CA3 PCs in awake mice and we propose mechanisms that underlie firing rate changes in CA3 during different brain states.

During theta, hippocampal pyramidal cells are engaged in encoding and recall of information. In a given environment, a proportion of CA3 PCs will be active while the majority are silent (Leutgeb, 2004). We found that theta was characterized by hyperpolarization of most CA3 PCs, in a manner which was closely time-locked to each theta event, suggesting a tight temporal coupling between the underlying mechanism and theta. This hyperpolarization was correlated with a decreased firing rate, consistent with the findings made with extracellular recording techniques (Leutgeb, 2004). Our data also show that these membrane potential dynamics are dependent on the brain state rather than the behavior of the animal, as they were seen both during run theta and rest theta.

During theta, we observed a decrease in the variance of the membrane potential in CA3 PCs regardless of the change in membrane potential, a slight decrease in cell input resistance, and a dependence on the initial membrane potential value, all of which point to a global increase in inhibition. However, we cannot pinpoint whether this apparent increase in inhibition is due to neuromodulation (Polack et al., 2013) or to synaptic inhibitory inputs, nor can we definitively exclude other mechanisms for hyperpolarization, such as a decrease in excitatory input. Overall, we have observed a robust phenomenon in intracellular dynamics that excludes most cells from putative ensembles of active neurons, thus ensuring a high signal-to-noise ratio in the population firing.

In contrast to our results, it has been reported that CA1 PCs recorded in similar conditions display no marked modulation of membrane potential during theta (Hulse et al., 2017). However, optical recording of CA1 PCs using voltage sensitive dyes shows a decrease in firing rate and in the power of subthreshold fluctuations during walking, a behavior associated with theta oscillations in the hippocampus (Adam et al., 2019). This modulation of CA1 PCs occurs together with increased activity of SST interneurons, possibly driven by the medial septum (Adam et al., 2019). It is possible that a similar mechanism is taking place in the CA3 network during theta and contributes to the modulations we report.

During LIA, the brain is in a low-arousal state and hippocampal pyramidal cells engage in offline replay of sequences associated with previously learned information (Buzsáki, 2015; Foster and Wilson, 2006). A detailed analysis of membrane potential fluctuations in CA1 PCs and in DG cells across transitions to LIA was recently made in awake head-fixed mice (Hulse et al., 2017). Most cells in both CA1 and DG depolarized and their membrane potential demonstrated increased variability. In a small proportion of the recorded cells, we observed similar features for CA3 PCs. While the population of CA3 PCs tended to depolarize, most cells did not do so consistently; instead they displayed different membrane potential modulation (hyperpolarization, depolarization or no change) across LIA events. We hypothesize that individual cells are activated during some LIA events but not others, which fits well with previous research showing that different SWRs activate specific ensembles (Ramirez-Villegas et al., 2015; Taxidis et al., 2015). The increased membrane potential variance in CA3 PCs and known depolarization in upstream DG suggests that CA3 PCs experience an increase in excitatory input during LIA, leading to an increase in firing rate in some cells even if the cells’ excitability is stable across brain states.

In this study, we show that during theta, CA3 PCs mostly behave in a stereotypical manner, characterized by a consistent hyperpolarization time-locked to the duration of the theta event. These conditions would favor a high signal-to-noise ratio which may allow a small subpopulation of CA3 PCs participating in a memory-linked neuronal ensemble to increase their downstream impact. In contrast, during LIA, CA3 PCs exhibit heterogeneous changes in the membrane potential, and only a subset consistently depolarize and increase firing. These results demonstrate that CA3 PCs show coordinated changes in membrane potential dynamics during theta which are consistent with a proposed role of increased inhibition leading to high-signal to noise ratio during a high-arousal encoding state. This work creates a strong basis for future modelling and physiological studies investigating cellular computations during brain-states associated with different hippocampal functions.

## Acknowledgements

We are grateful to Ania Gonçalves for help in confocal imaging and Mario Carta, Lisa Roux, Xavier Leinekugel, Ruth Betterton, and Catherine Marneffe for useful comments on the manuscript and fruitful discussions. This study was supported by the Centre National de la Recherche Scientifique, the European Commission (EIF Fellowship awarded to A.K.; 702037), the ANR (grant Hippencode #14-CE13-0015), and the Fondation pour la Recherche Médicale (to M.M.; FDT20170437372).

## Author Contributions

Conceptualization, M.M., A.K. and C.M.; Methodology, M.M., A.K.; Investigation, M.M., A.K.; Formal Analysis A.K. and M.M.; Writing - Original Draft M.M., A.K. and C.M.; Visualization M.M. and A.K.; Supervision C.M.; Project administration C.M.; Funding acquisition M.M., A.K. and C.M.

---

**Table S1.** Cell properties and state-associated modulations in Vm.

| Cell ID | Mouse ID | Total Recording Time (s) | Access Resistance (MΩ) | Input Resistance (MΩ) | Capacitance (pF) | Number of Theta Events |  |  | Theta Classification | Number of LIA Events |  |  | LIA Classification |
| --- | --- | --- | --- | --- | --- | --- | --- | --- | --- | --- | --- | --- | --- |
|  |  |  |  |  |  | hyp | no | dep |  | hyp | no | dep |  |
| 1 | 1 | 1080 | - | - | - | 4 | 5 | 1 |  | 11 | 11 | 12 |  |
| 2 | 2 | 600 | - | - | - | 14 | 11 | 17 |  | 1 | 3 | 5 |  |
| 3 | 3 | 450 | 54.7 | 163.1 | 477.4 | 9 | 5 | 7 |  | 3 | 2 | 6 |  |
| 4 | 4 | 300 | 70.2 | 172.3 | 100.7 | 11 | 6 | 1 | hyperpolarize | 1 | 3 | 4 |  |
| 5 | 5 | 300 | 33.9 | 62.8 | 682.0 | 21 | 1 | 0 | hyperpolarize | 0 | 0 | 1 |  |
| 6 | 6 | 40 | 72.1 | 123.5 | 168.8 | 1 | 2 | 0 |  | 0 | 0 | 0 | - |
| 7 | 7 | 150 | 36.1 | 91.5 | 193.3 | 0 | 1 | 1 |  | 2 | 2 | 1 |  |
| 8 | 7 | 90 | 48.7 | 197.4 | 186.2 | 1 | 1 | 0 | hyperpolarize | 0 | 1 | 1 |  |
| 9 | 8 | 1065 | 61.3 | 124.4 | 123.5 | 33 | 6 | 10 |  | 4 | 11 | 26 |  |
| 10 | 9 | 490 | 24.3 | 108.4 | 243.1 | 7 | 0 | 1 | hyperpolarize | 8 | 7 | 8 |  |
| 11 | 10 | 1395 | 16.3 | 145.6 | 217.2 | 25 | 56 | 13 | hyperpolarize | 5 | 6 | 10 |  |
| 12 | 10 | 250 | 31.2 | 122.8 | 220.9 | 16 | 1 | 1 | hyperpolarize | 1 | 2 | 1 |  |
| 13 | 11 | 1105 | 76.8 | 260.0 | 156.5 | 27 | 23 | 10 | hyperpolarize | 6 | 21 | 18 | depolarize |
| 14 | 11 | 475 | 27.5 | 210.4 | 274.2 | 18 | 1 | 0 | hyperpolarize | 2 | 2 | 7 | depolarize |
| 15 | 12 | 390 | 61.7 | 220.9 | 213.0 | 19 | 5 | 1 | hyperpolarize | 1 | 2 | 2 |  |
| 16 | 12 | 345 | 56.8 | 174.3 | 211.6 | 3 | 3 | 5 |  | 3 | 4 | 4 |  |
| 17 | 12 | 510 | 34.6 | 78.7 | - | 5 | 4 | 1 |  | 6 | 7 | 11 |  |
| 18 | 13 | 375 | 83.3 | 347.7 | 144.4 | 4 | 2 | 1 |  | 0 | 7 | 9 | depolarize |
| 19 | 14 | 660 | 60.3 | 313.2 | 110.2 | 17 | 9 | 5 | hyperpolarize | 2 | 6 | 10 | depolarize |
| 20 | 15 | 250 | 49.5 | 172.7 | 82.4 | 5 | 3 | 4 |  | 1 | 4 | 4 |  |
| 21 | 15 | 200 | 65.6 | 158.3 | 119.0 | 4 | 3 | 7 |  | 0 | 0 | 1 |  |
| 22 | 16 | 250 | 84.7 | 261.7 | 153.7 | 8 | 3 | 1 | hyperpolarize | 0 | 5 | 2 | depolarize |
| 23 | 16 | 290 | 66.8 | 185.2 | 172.4 | 13 | 4 | 2 | hyperpolarize | 2 | 2 | 4 |  |
| 24 | 16 | 300 | 55.3 | 118.2 | 94.5 | 5 | 8 | 4 |  | 2 | 1 | 1 |  |
| 25 | 17 | 300 | 29.6 | 188.0 | 177.6 | 11 | 9 | 8 |  | 0 | 4 | 0 |  |
| 26 | 17 | 300 | 58.9 | 179.4 | 109.0 | 7 | 9 | 11 | depolarize | 0 | 2 | 2 |  |
| 27 | 18 | 300 | 47.1 | 146.7 | 133.2 | 18 | 3 | 0 | hyperpolarize | 1 | 4 | 6 | depolarize |
| 28 | 18 | 300 | 48.7 | 117.9 | 157.9 | 9 | 5 | 3 | hyperpolarize | 1 | 8 | 3 |  |
| 29 | 19 | 290 | 55.1 | 141.9 | 113.9 | 5 | 1 | 1 | hyperpolarize | 4 | 6 | 1 |  |
| 30 | 19 | 150 | 36.0 | 95.4 | 147.2 | 4 | 2 | 3 |  | 2 | 2 | 1 |  |
| 31 | 19 | 80 | 40.8 | 99.3 | 137.0 | 4 | 0 | 0 | hyperpolarize | 0 | 1 | 1 |  |
| 32 | 20 | 200 | 41.0 | 121.1 | 123.2 | 6 | 3 | 4 |  | 0 | 1 | 5 | depolarize |
| 33 | 21 | 150 | 37.3 | 183.2 | 313.4 | 9 | 0 | 0 | hyperpolarize | 0 | 0 | 3 | depolarize |
| mean ± std |  | 407.0 ± 315.6 | 50.5 ± 17.3 | 164.1 ± 64.8 | 191.9 ± 119.5 | - | - | - | - | - | - | - | - |

**Table S2.** Cell spiking properties.

| Cell ID | Mouse ID | Average Firing Rate (Hz) | Spikes Occurring Within Complex Spikes (%) | Average Frequency of Complex Spikes (Hz) | Average ISI Within Complex Spikes (ms) | Average Number of Spikes in a Complex Spike | Number of Spikes Used in Threshold Analysis | Average Full Width at Half-Max (ms) | Average Maximum Spike Slope (V/s) |
| --- | --- | --- | --- | --- | --- | --- | --- | --- | --- |
| 1 | 1 | 0.6 | 69 | 0.04 | 7.2 | 12 | 200 | 0.9 | 77.8 |
| 2 | 2 | 0.1 | 69 | 0.01 | 12.5 | 18 | 21 | 0.9 | 165.7 |
| 3 | 3 | 0.9 | 52 | 0.08 | 6.2 | 6 | 184 | 1.2 | 70.7 |
| 4 | 4 | 0.4 | 97 | 0.04 | 4.5 | 9 | 7 | 0.8 | 132.1 |
| 5 | 5 | 5.3 | 41 | 0.49 | 5.5 | 4 | 708 | 1.2 | 64.0 |
| 6 | 6 | 3.6 | 83 | 0.40 | 4.4 | 7 | 32 | 1.0 | 69.7 |
| 7 | 7 | 6.5 | 29 | 0.32 | 7.6 | 6 | 529 | 0.8 | 150.2 |
| 8 | 7 | 4.6 | 82 | 0.50 | 5.3 | 8 | 99 | 1.0 | 106.1 |
| 9 | 8 | 3.4 | 61 | 0.29 | 6.7 | 7 | 1226 | 1.1 | 60.8 |
| 10 | 9 | 1.9 | 61 | 0.15 | 8.0 | 8 | 353 | 1.2 | 89.1 |
| 11 | 10 | 0.1 | 71 | 0.01 | 6.2 | 8 | 40 | 1.0 | 122.9 |
| 12 | 10 | 2.1 | 75 | 0.18 | 4.7 | 9 | 135 | 1.0 | 79.6 |
| 13 | 11 | 0.1 | 46 | 0.01 | 7.9 | 5 | 31 | 1.4 | 74.0 |
| 14 | 11 | 0.3 | 90 | 0.03 | 6.9 | 7 | 19 | 0.8 | 239.4 |
| 15 | 12 | 1.4 | 80 | 0.16 | 4.8 | 7 | 126 | 1.1 | 82.5 |
| 16 | 12 | 0.3 | 36 | 0.03 | 8.3 | 4 | 58 | 1.0 | 115.5 |
| 17 | 12 | 0.4 | 98 | 0.02 | 6.4 | 23 | 10 | 1.3 | 62.6 |
| 18 | 13 | 0.2 | 56 | 0.02 | 7.3 | 6 | 21 | 1.2 | 73.1 |
| 19 | 14 | 0.1 | 76 | 0.01 | 7.4 | 10 | 13 | 1.1 | 82.3 |
| 20 | 15 | 0.1 | 0 | 0.00 | - | - | 17 | 1.3 | 54.5 |
| 21 | 15 | 1.3 | 50 | 0.10 | 6.2 | 7 | 110 | 1.1 | 76.6 |
| 22 | 16 | 0.0 | - | - | - | - | - | - | - |
| 23 | 16 | 0.3 | 58 | 0.02 | 8.9 | 9 | 24 | 1.1 | 92.1 |
| 24 | 16 | 1.5 | 94 | 0.10 | 8.1 | 13 | 47 | 1.4 | 62.7 |
| 25 | 17 | 5.6 | 10 | 0.12 | 10.3 | 5 | 1025 | 0.8 | 177.0 |
| 26 | 17 | 2.5 | 46 | 0.17 | 7.3 | 7 | 284 | 1.2 | 64.6 |
| 27 | 18 | 0.5 | 83 | 0.05 | 4.8 | 8 | 31 | 1.0 | 101.2 |
| 28 | 18 | 3.4 | 74 | 0.29 | 6.5 | 9 | 240 | 0.8 | 137.1 |
| 29 | 19 | 4.5 | 45 | 0.34 | 7.7 | 6 | 554 | 1.2 | 83.4 |
| 30 | 19 | 0.0 | - | - | - | - | - | - | - |
| 31 | 19 | 4.4 | 78 | 0.44 | 7.9 | 8 | 77 | 1.0 | 121.4 |
| 32 | 20 | 0.9 | 97 | 0.07 | 4.4 | 12 | 17 | 1.1 | 85.2 |
| 33 | 21 | 2.4 | 36 | 0.18 | 6.2 | 5 | 169 | 1.0 | 89.1 |
| *mean ± std* |  | *1.8 ± 1.9* | *63 ± 25* | *0.14 ± 0.15* | *6.9 ± 1.8* | *8 ± 4* | *194 ± 293* | *1.1 ± 0.2* | *98.8 ± 40.5* |

## Methods

All experiments were approved by the Ethical Committee #50 and the French Ministry for Education and Research.

### Animals

C57Bl6/j male mice were obtained from Charles River and cared for according to the regulations of the University of Bordeaux/CNRS Animal Care and Use Committee. Animals were housed with their littermates with *ad libitum* access to food and water. Cages were kept in a temperature-regulated room on a 12 h light/dark cycle. Electrophysiological recordings were performed on P45-to-P71 mice.

### Surgery

#### Headbar implantation

Custom-made stainless steel head bars (35x3x3 mm) were affixed to the skulls of 4-to-5-week-old mice using Superbond (Phymep) and dental acrylic (Dentalon, Phymep) while under isoflurane anesthesia. The skull underneath the head implant opening was covered with superbond and future locations for the craniotomies above CA3 were marked using stereotactic coordinates (2.0 caudal, 2.0 lateral from bregma).

#### Craniotomy

The day of recording, animals were anesthetized using isoflurane and a craniotomy (∼1.5 mm diameter) was drilled at previously marked location. The dura was left intact and craniotomy was covered with silicone elastomer (Kwik-cast, World Precision Instruments). Mice were placed in a recovery cage for at least 1.5 hours before recording. Mice were typically recorded over two consecutive days.

### Training

After headbar implantation, mice were left 2-3 days for recovery, until they started to gain weight again. On the first day of training, mice were allowed to freely explore the wheel for 15 min without being headfixed. During the following 4 days, mice were progressively headfixed (10 min, 30 min, 60 min, and 120 min). Thereafter, mice were headfixed 120 min each day until they habituated and were alternating between running and stopping without showing signs of excessive stress (freezing, escape attempts, struggling, excessive excretion, vocalization). Typically, mice habituated in 2 to 10 days. During the final two days of training, mice were headfixed on the wheel used for the recordings and habituated to the ambient light and noises that would occur during an experiment.

### Recordings

#### Whole-cell recordings

Before attempting whole-cell patching, the depth of the pyramidal cell layer was detected using an extracellular electrode. Extracellular electrodes were glass patch electrodes (World Precision Instruments) prepared with a vertical puller PC-10 (Narishige) and then broken so that the resistance when filled with standard Ringer’s solution (in mM: 135 NaCl, 5.4 KCl, 5 HEPES, 1.8 CaCl2, and 1 MgCl2) was 0.8–1.5 MΩ. The extracellular electrode used to determine the pyramidal cell depth for whole-cell recording was mounted vertically on a micromanipulator (Scientifica). As the electrode was advanced through the cortex, the CA1 pyramidal cell layer was identified when multi-unit activity could be detected and the amplitude of ripples increased, typically 1.3–1.5 mm below the surface of the brain. The polarity of the sharp waves inversed below the CA1 pyramidal layer, and the electrode was advanced further down the hippocampus until the sharp waves changed polarity again and multi-unit activity was detected, indicative of the CA3 pyramidal layer. The extracellular electrode to be used for LFP recordings during whole-cell patching was mounted on a second micromanipulator (Luigs and Neumann) at a 22.5° angle relative to the vertical axis, and advanced through the same craniotomy at a more medial and caudal position. This extracellular electrode was advanced until detection of the pyramidal cell layer but not through it. The two LFPs were recorded for visual confirmation of a high correlation between the two signals (phase of theta, coincidence of ripples, multi-unit activity), indicative of a short distance between the two recording sites. Whole-cell patch electrodes (3.5-5 MΩ) were prepared beforehand and were filled with a solution containing (in mM) 134 K-methanesulfonate, 2 KCl, 10 HEPES, 10 EGTA, 2 MgCl2, and 2 Na-ATP, at 290–300 mOsm and pH 7.2–7.3 adjusted with KOH. The intracellular solution was supplemented with 0.2% (w/v) biocytin for post hoc cellular identification and morphological reconstruction. Whole-cell patch-clamp recordings were achieved using a standard blind-patch approach as previously described (Lee et al., 2009; Margrie et al., 2002). Access resistance and membrane capacitance of the cells were monitored on-line with the membrane test feature of the acquisition software and analyzed off-line. The mean access resistance was 52.4 ± 2.5 MΩ. Membrane potential was not corrected for liquid junction potential which was estimated to be approximatively 11 mV (LJP calculator, Clampex). Recordings were discarded when the access resistance exceeded 100 MΩ or when the action potential peak dropped below 0 mV.

### Data acquisition

#### Electrophysiology

All recordings were made in the current-clamp configuration. Recordings were obtained with a Multiclamp 700B amplifier connected to the Digidata 1440A system. Data were acquired with pClamp 10 (Molecular Devices), digitized at 10 kHz, filtered at 3 kHz, and analyzed off-line with Clampfit 10.4 (Molecular Devices) and Python. To measure input resistance, we injected trains of current pulses (-100 pA, 100 ms) in a separate set of recordings (n = 13).

#### Pupil recording

Pupil diameter was measured for a subset of recordings (n=11/33). To measure pupil diameter, the mouse was illuminated with an infrared (850 nm) battery-operated light (Maketheone IR Torch 850NM) and imaged with a camera (DALSA GENIE NANO-M1280-NIR) at 10 Hz.

#### Locomotion

Online running behavior and the animal’s velocity during the recording session was measured using Poly Wheel software (Imetronic, France) and acquired at 32 Hz. Locomotion was detected as IR-sensor-crossing by the animal. 16 sensors were located on a wheel of 20 cm diameter.

### Histology

To recover the morphology of neurons filled with biocytin during *in vivo* whole-cell patch-clamp recordings, brains were removed after the completion of electrophysiology experiments and were fixed for 1 week at 4°C in a solution containing 4% PFA in 1X PBS, pH7.4. Coronal slices (100 µm thick) were incubated overnight at 4°C in 1:1000 Alexa Fluor 488 conjugated in the same solution. Images were acquired using a confocal microscope (TCS SP8, Leica) and oil objective HC PL APO 40x/1.30.

### Analysis

#### LFP

LFP signal was downsampled 10 times to 2000 Hz and filtered using a 4-pole butterworth band-pass filter between 0.2 Hz and 100 Hz. After filtering, LFP was again downsampled to 200 Hz. The spectrogram of the LFP was calculated using consecutive fast Fourier transforms within ∼2 second boxcar windows with 99% overlap. Theta/delta ratio was found by dividing the average power in the theta range (6-9 Hz) by the average power in the delta range (0.5-3.5 Hz). This ratio was normalized within sessions by taking the z-score. For the total power, the spectrogram was z-scored within frequency bands, and averaged between 0.3 and 80 Hz.

#### Brain state classification

Theta events were identified as periods with a theta/delta power ratio above the mean. To be counted as theta events, theta bouts had to be longer than 1 s and have a power ratio higher than 1 sd above the mean at least once. If two theta bouts happened less than 1 s apart, they were merged into one single theta period. Theta events were considered rest theta when the running speed on the wheel was zero for the entire duration of the event; theta events that overlapped with any period of nonzero speed were considered run theta. LIA events were identified as periods with a total LFP power above the mean. To be counted as LIA events, LIA bouts had to be longer than 2 s and have a total power higher than 0.25 sd above the mean at least once. If two LIA bouts happened less than 0.5 s apart, they were merged into one single LIA period.

#### Membrane potential

Analysis of membrane potential changes was performed on membrane potential traces after spike removal. Spikes were detected when the derivative of the membrane potential (dV/dt) exceeded 5 mV/ms. To keep a conservative estimate of average membrane potential, spikes and their underlying depolarizations such as plateau potentials were removed. The spike-removed membrane potential trace was then smoothed by taking the mean over a sliding window of 2 s, and the smoothed membrane potential was downsampled to 20 Hz. The residual membrane potential trace was obtained by subtracting the smoothed membrane potential from the spike-removed membrane potential. From this residual membrane potential, the variance was calculated over a sliding a window of 1 s. The variance trace was then smoothed by taking the mean over a sliding window of 2 s and downsampled to 20 Hz. For the measurements of membrane potential variance, we found that it was important to remove the subthreshold components of spikes using the procedure described above. If the subthreshold components of the complex spikes were left in the trace (as described in the CS analysis below), the histograms of membrane potential would be skewed to depolarized values and bias the variance measurements.

#### Input resistance

Input resistance was calculated using the voltage response to 100 ms steps of -100 pA current. The average voltage during the 250 ms before the current pulse was subtracted from the 250 ms before the end of the current pulse, and the result was divided by the amplitude of the current pulse. Input resistance measurements were discarded when spikes were detected occurring in a window of 750 ms centered around either sample period.

#### Detection of complex spikes

The time stamp and threshold potential for each spike was taken as the time and membrane potential when the first derivative (dV/dt) first surpassed 5 V/s (Epsztein et al., 2011) The end of each spike was detected as the first local minimum after the peak of the spike. A subthreshold potential trace was created by removing the spikes and linearly interpolating between the start and end of the spikes. This method preserved the subthreshold depolarizations characteristic of complex spikes. Complex spike initiation was detected as any group of three or more spikes that had an isi of 20 ms or less. The start time and voltage of the complex spike was defined as that of the threshold of the first spike in the burst. The end of the complex spike was defined as the first local minimum after the subthreshold potential returned to within 3 mV of the starting voltage of the complex spike. Any spikes that occurred between the start and the end of the subthreshold component of the complex spike were considered part of that complex spike, regardless of their inter-spike interval.

#### Spike threshold

The time stamp and threshold potential for each spike was taken as the time and membrane potential when the first derivative (dV/dt) first surpassed 5 V/s, as above. It is known that the spike threshold increases for spikes within a CS after the first spike (Kandel et al., 1961)so care was taken to include in the threshold analysis only isolated spikes, or the first spike in a complex spike. Spikes that had a prior inter-spike interval of less than 50 ms were eliminated from the analysis. In addition, spikes that were in a CS, but not the first spike, were also eliminated. Spikelets were also eliminated by discarding spikes that occurred outside of CS and did not reach peak amplitude of at least -10 mV. The distance to spike threshold for each spike was calculated by subtracting the spike threshold from the baseline membrane potential over a specified period prior to the spike. The baseline membrane potential was defined as the mode of the voltages after spike removal (replacing spikes and their subthreshold components with nans instead of linearly-interpolated values), smoothing over windows of 2ms, downsampling to 500 Hz, and rounding all values to the nearest 0.1 mV. For the relative spike threshold, this baseline was taken in the 50-300 ms window prior to the spike. For the pre-spike depolarization (PSD), this baseline was taken from the smoothed, downsampled membrane potential 20 ms prior to the spike.

#### Detection of hyperpolarizations and depolarizations

To determine whether a particular event was associated with a significant change in membrane potential, 2 two-second windows of the 20 Hz downsampled membrane potential around the start of the event were compared with a Welch’s t-test. For theta, the comparison was made between –2.5 to –0.5 and 0.5 to 2.5 seconds with respect to the start of the event; for LIA, the windows were –4 to –2 and –1 to 1 second. These windows were chosen according to the times of maximal changes in the grand average of event-triggered membrane potential traces. Events with p-values less than 0.05 were considered to be hyper- or depolarizing depending on the direction of the change in membrane potential. To detect hyperpolarizations and depolarizations not associated with state onsets, the same method was used, testing all times in the recording for a significant change in membrane potential over the specified windows. Then the detected hyperpolarization and depolarization times were compared with those of theta and LIA onsets to separate those associated with either state from those associated with no state.

#### Pupil diameter

To measure pupil diameter, frames were extracted from the video file. A Circle Hough Transform (CHT) was performed to detect circular objects in each individual frames. The radius of the detected circles was extracted automatically. Pupil diameter data was included after control of correct pupil detection. The resulting pupil diameter trace was smoothed with a 4-pole butterworth low-pass filter (0.5 Hz cutoff).

### Statistics

Unless otherwise noted, data were expressed as the median +/- the mean absolute derivation about the median, which was calculated by subtracting each data point from the group’s median, taking the absolute value, summating, and then dividing by the number of data points. These measures of central tendency and variation were used because many distributions were found not to be normal. When comparing 2 independent groups, a bootstrapping method was used. Briefly, data from both groups were randomly resampled with replacement to form two surrogate groups. This was repeated 1,000 times, and if the difference in medians was more extreme than 95% of the surrogate trials, the groups were considered significantly different. For comparisons of 3 or more groups, a Kruskal-Wallis test was performed to determine the presence of any significantly different group before performing the 2-group bootstrap for each group pair. A different bootstrap test was used when comparing two non-independent groups. In this test, the difference between paired data points was randomly determined to be either positive or negative 1,000 times. If the mean of the differences in the actual data set was more extreme than 95% of those in the shuffled data set, the paired groups were considered to have a significant difference. For comparative scatter plots where variables during different states were compared, a Mann-Whitney U test was used to determine whether there was a significant difference between the states for that particular cell (filled circles in scatter plots). When determining whether a value was significantly different from zero, a bootstrapping method was used. The distribution was demeaned and then resampled with replacement 10,000 times. If the mean of the original data was more extreme than 95% of the resampled means, it was considered significantly different from zero.

To determine whether a particular event was associated with a significant change in membrane potential, 2 two-second windows of the 20 Hz downsampled membrane potential around the start of the event were compared with a Welch’s t-test. For theta, the comparison was made between –2.5 to –0.5 and 0.5 to 2.5 seconds with respect to the start of the event; for LIA, the windows were –4 to –2 and –1 to 1 second. Events with p-values less than 0.05 were considered to be hyper- or depolarizing depending on the direction of the change in membrane potential. To determine whether the proportions of hyperpolarizing, depolarizing and no change events could be due to chance, a bootstrapping method was used. For every recording, the actual event times were replaced with randomly-sampled times, and the same method as previously described was used to determine whether each random timepoint was associated with a hyperpolarization, depolarization, or no change. This was repeated 1,000 times to create a distribution of proportions. Proportions were considered significantly different from chance if they were either greater or smaller than 95% of the randomly-sampled data sets.

To determine whether a cell was consistently changed membrane potential at the start of a particular type of event, we used a bootstrapping method. Over a single recording session, the membrane potential and actual event times were shifted in time with respect to one another 6,000 times in increments of 50 ms. For each shift, the average change in membrane potential associated with event onset was calculated for the cell. Cells were considered significantly hyper- or depolarizing when the average change in membrane potential was more extreme than 95% of the shuffled data.

---

**Figure S1.**
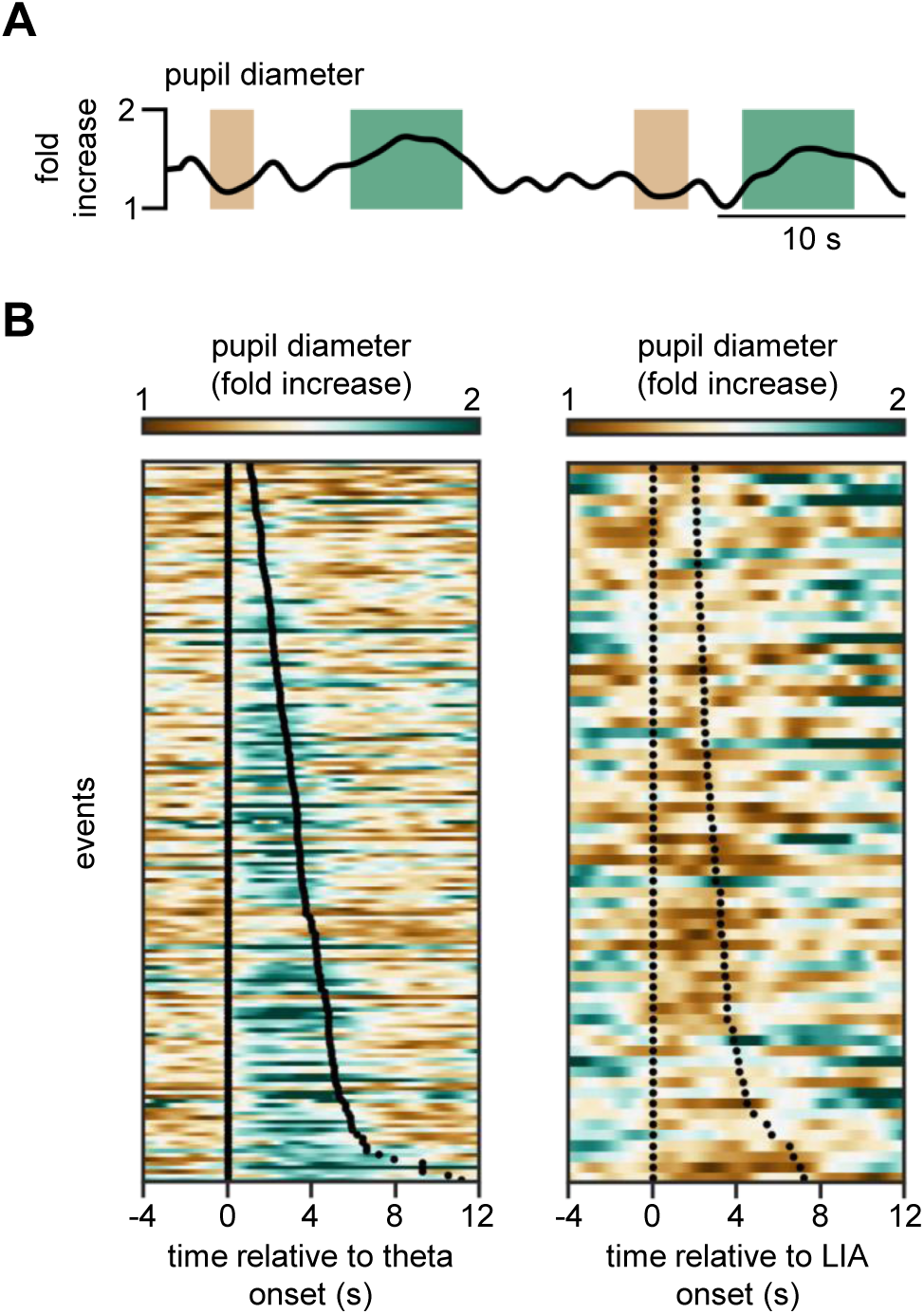
Pupil dilation during theta and constriction during LIA. (A) Example trace of pupil dilation showing dilations during theta (green) and constriction during LIA (brown). (B) Color plots showing pupil diameter triggered by transition to theta (left, n = 174 events) and LIA (right, n = 69 events) from the subset of recordings with pupil diameter tracking (n = 12 cells). Events are organized according to the length of the brain state, with onset and offset represented by the two black dotted lines.

**Figure S2.**
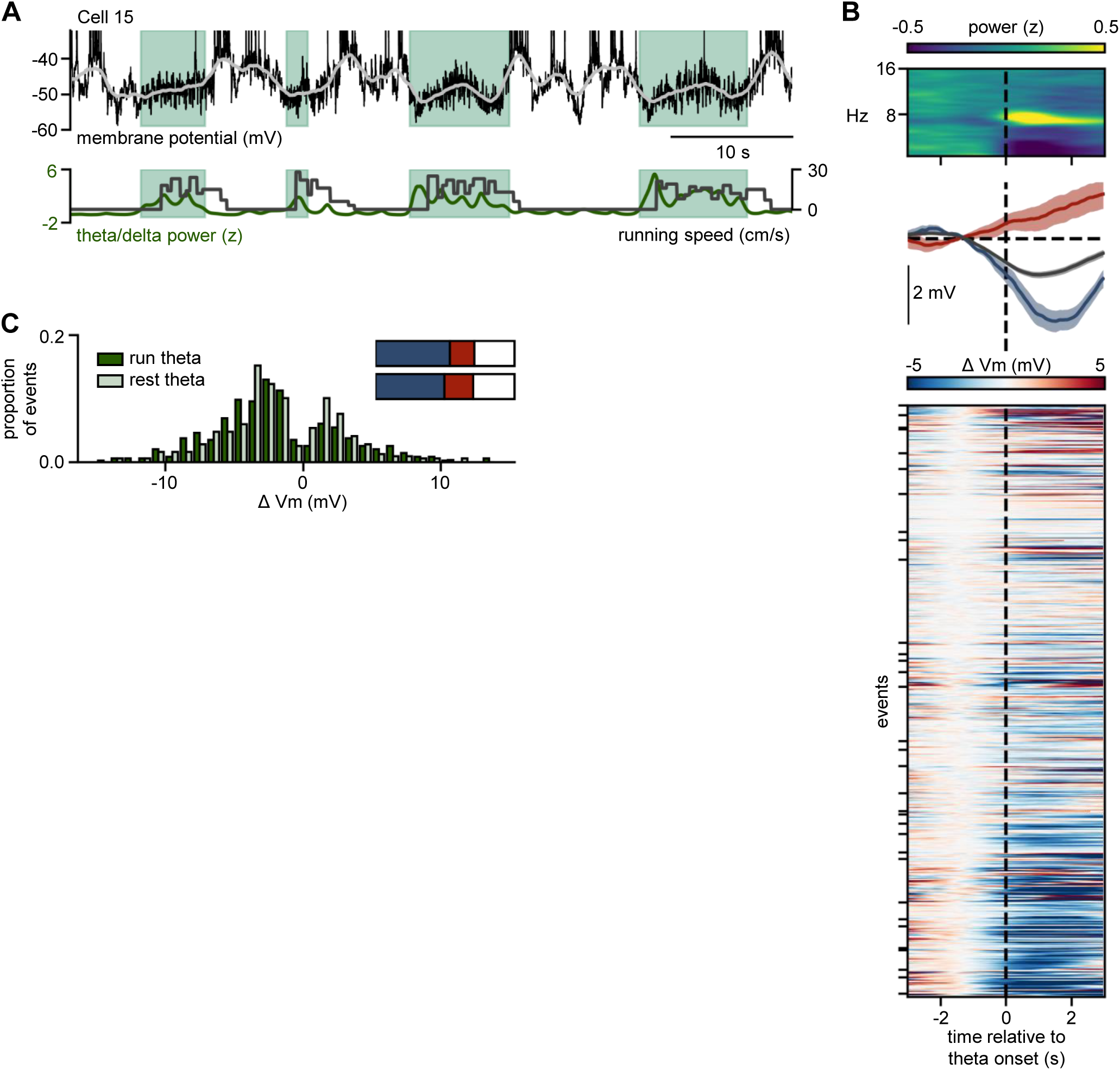
CA3 pyramidal cells hyperpolarize during theta. (A) Example whole-cell recordings from a CA3 PC (top), with simultaneous theta/delta power ratio of the nearby LFP and the running speed of the mouse (bottom). Grey trace superimposed on the raw Vm is the smoothed Vm after spike removal, and green shading represents theta events. Spikes are truncated in the raw membrane potential traces. (B) Top: Spectrogram of the average LFP (z-scored within frequency bands) triggered by transitions to theta (time = 0, dotted line) showing an increase of power in the theta band (6-9 Hz). Middle: Average Vm traces during theta onset of an example hyperpolarizing (blue) and depolarizing (red) cell, as well as the grand average over all events (grey) (n = 668, shaded areas represent ± sem). Bottom: Color plot showing the Vm of CA3 PCs triggered by transition to theta normalized to 2.5 to 0.5 s before start of theta. Blue represents Vm hyperpolarization, red Vm depolarization. Each cell is represented in between black ticks. Cells are ordered depending on their average Vm modulation during theta, with depolarizing cells on top and hyperpolarizing cells at the bottom of the plot. Events within each cell are ordered by the time in which they occurred in the recording. (C) Comparison of Vm modulation during theta for run [n = 350] and rest [n = 318] events (bin size = 1 mV). Inset: bar charts of percent event type (hyperpolarizing, depolarizing, or no change) for run (top) and rest (bottom) theta events. All types of events occurred at a similar frequency during run and rest theta (p = 0.739 for hyperpolarizing, p = 0.201 for depolarizing, p = 0.383 for no change).

**Figure S3.**
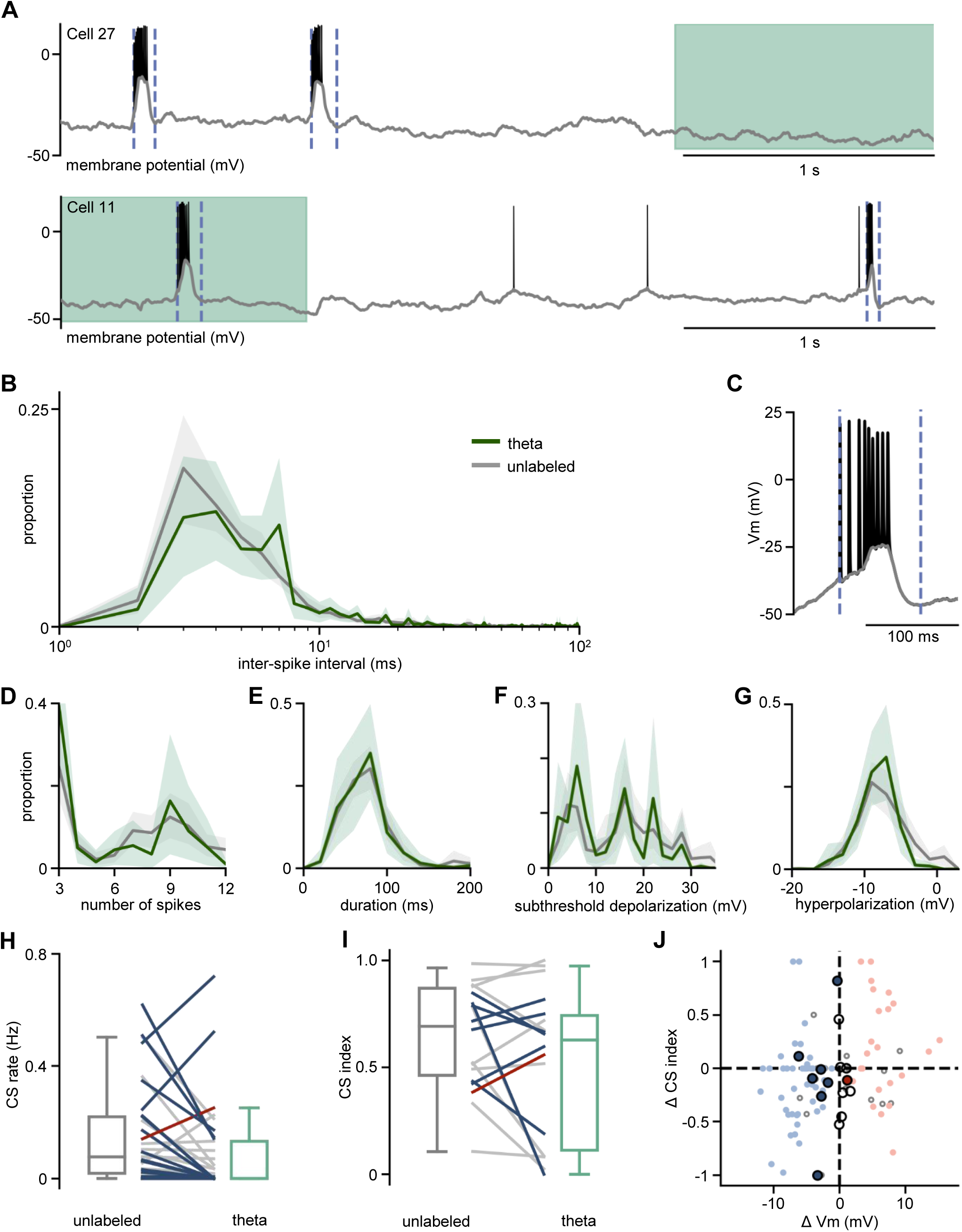
State-associated change in Vm does not influence complex spikes. (A) Example membrane potential traces from two different theta hyperpolarizing cells. Raw traces (black) are superimposed with subthreshold trace (grey). Blue vertical lines represent the beginning and end of complex spikes. (B) Proportion of the inter-spike interval for each state, averaged over cells. Distribution is similar across states (8.9 ± 14.1 ms [n = 30 cells] for unlabeled, 12.0 ± 44.4 ms [n = 15 cells] for theta, paired test over cells: p = 0.869). Only cells with at least 10 spikes in the state were included in the corresponding statistics. (C) Example trace showing a representative CS. Blue dotted lines represent beginning and end of the CS used to calculate CS duration and amplitude of subthreshold depolarization and after hyperpolarization. Grey line represents the subthreshold membrane potential. (D) – (G) CS properties for each state; distributions are separated by state and averaged over cells. (D) Number of spikes per CS was not different between theta and unlabeled (7 ± 3 spikes [n = 29 cells] for unlabeled, 7 ± 3 spikes [n = 16 cells] for theta, paired test over cells: p = 0.562). (E) CS duration was not different between theta and unlabeled (82.3 ± 18.0 ms [n = 29 cells] for unlabeled, 82.8 ± 16.0 ms [n = 16 cells] for theta, paired test over cells: p = 0.786). (F) Subthreshold depolarization of the CS was not different between theta and unlabeled (16.8 ± 6.0 mV [n = 29 cells] for unlabeled, 13.0 ± 5.6 mV [n = 16 cells] for theta, paired test over cells: p = 0.117). (G) Cells had deeper hyperpolarizations after CS during theta than during unlabeled states (-6.6 ± 1.8 mV [n = 29 cells] for unlabeled, -7.7 ± 1.3 mV [n = 16 cells] for theta, p = 0.016). (H) CS rate during theta and unlabeled states. Colored lines represent cells with significant Vm modulation during each state. The CS rate was lower during theta than during unlabeled (0.0 ± 0.1 Hz [n = 33 cells] for theta, 0.1 ± 0.12 Hz [n = 33 cells] for unlabeled, p = 0.013). (I) Same as in (H), but for CS index, defined as the proportion of spikes that occur within CSs. The CS index does not significantly differ between states for cells in which a CS index could be calculated (0.6 ± 0.3 [n = 18 cells] for theta, 0.7 ± 0.2 [n = 31 cells] for unlabeled, p = 0.055). (J) Change in CS index versus change in Vm during theta. Large dots represent average changes for a single cell (n = 15 cells). Small dots represent average for a single event (n = 87 events). Filled dots represent significant Vm modulation during theta, color represents the direction of modulation.

**Figure S4.**
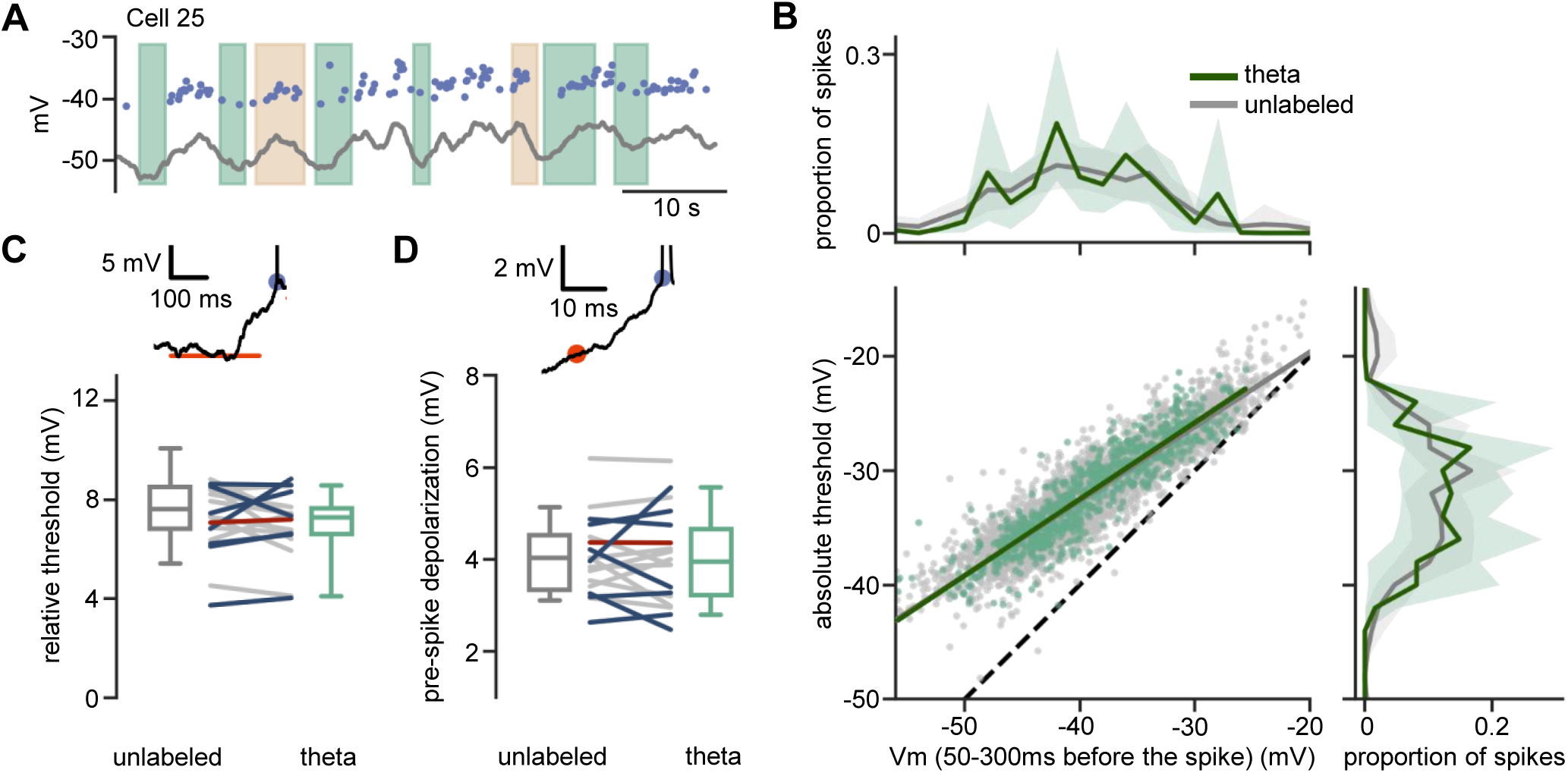
Dynamic changes in spike threshold relative to membrane potential during theta. (A) Example trace showing smoothed Vm (grey line) and spike thresholds (blue dots). Shaded areas represent states. (B) Bottom left: Absolute spike threshold vs baseline Vm value 50-300 ms before the spike. Dots represent single spikes, color represents the state in which the spike occurs. The slopes and intercepts of the linear regressions were not significantly different between states (y = 0.65x – 6.6 for unlabeled [n = 3705 spikes], y = 0.67x – 5.7 for theta [n = 853 spikes], change in slope paired test over cells: p = 0.055, change in intercept paired test over cells: p = 0.055). Top: Histograms of the baseline Vm prior to spikes, separated by state and averaged over cells (-38.0 ± 4.9 mV [n = 31 cells] for unlabeled, -38.9 ± 4.4 mV [n = 18 cells] for theta, paired test over cells: p = 0.572). Right: Histograms of the spike threshold, separated by states and averaged over cells (-30.9 ± 4.0 mV [n = 31 cells] for unlabeled, -31.2 ± 3.5 mV [n = 18 cells] for theta, paired test over cells: p = 0.709). (C) Top: example spike (truncated) showing the absolute spike threshold (blue dot) and the Vm 500-300 ms before the spike (orange line). Bottom: Relative threshold (difference between absolute spike threshold and baseline Vm) for each cell during theta and unlabeled states. Colored lines represent cells that significantly hyperpolarize (blue) or depolarize (red) during theta. There was no significant difference in the relative threshold over states (7.3 ± 1.0 mV [n = 18 cells] for theta, 7.6 ± 1.3 mV [n = 31 cells] for unlabeled, p = 0.246). (D) Top: example spike (truncated) showing the absolute spike threshold (blue dot) and the Vm 20 ms before the spike (orange dot). Bottom: Same as (C) but for the pre-spike depolarization. There was no significant difference in the pre-spike depolarization over states (3.9 ± 0.8 mV [n = 18 cells] for theta, 4.0 ± 0.6 mV [n = 31 cells] for unlabeled, p = 0.444).

**Figure S5.**
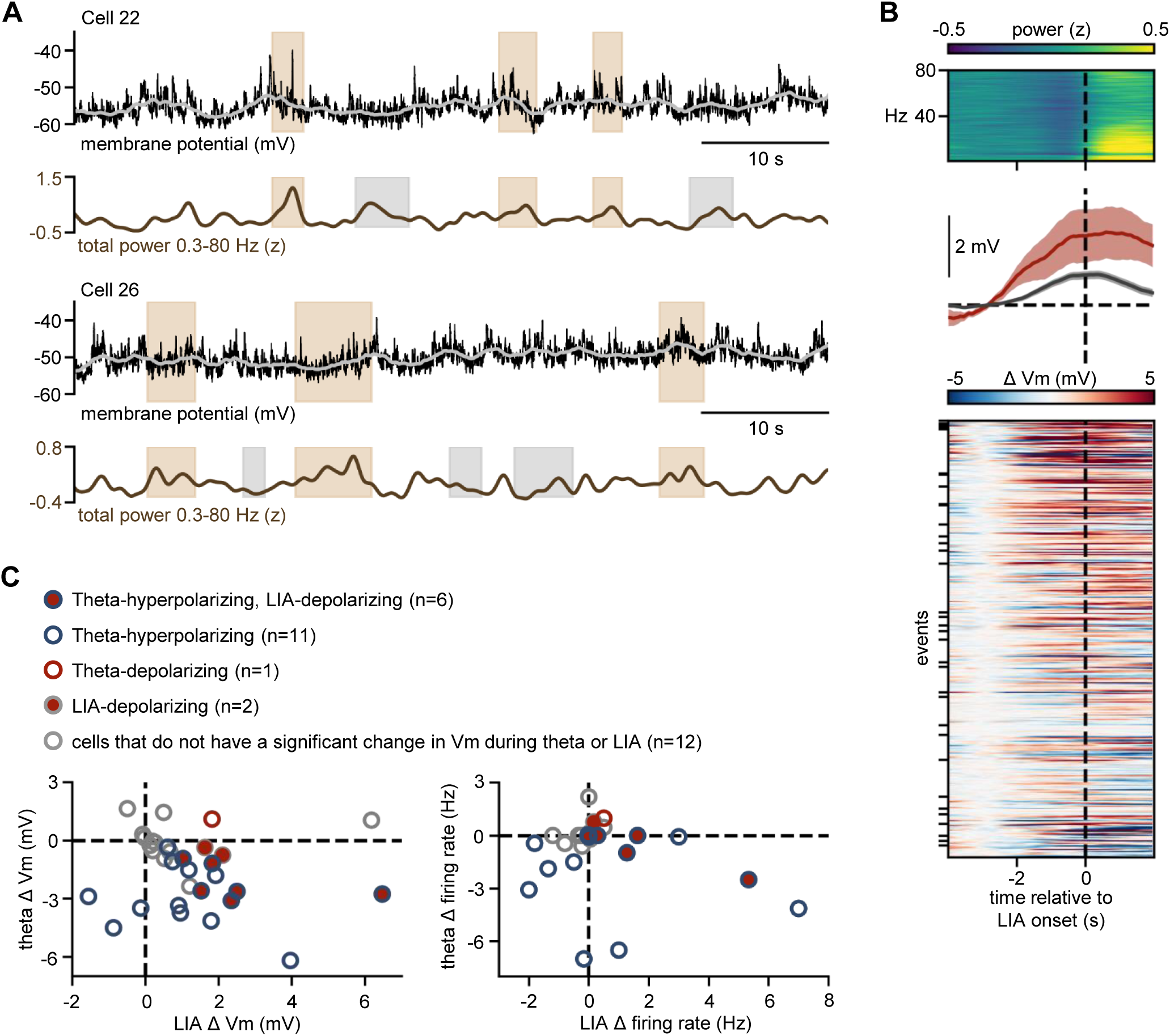
CA3 pyramidal cells have heterogeneous changes in membrane potential during LIA. (A) Example whole-cell recordings from two different CA3 PCs (top), with simultaneous total power of the nearby LFP (bottom). Grey trace superimposed on the raw Vm is the smoothed Vm after spike removal, and brown shading represents LIA events. Spikes are truncated in the raw membrane potential traces. (B) Top: Spectrogram of the average LFP (z-scored within frequency bands) triggered by transitions to LIA (time = 0, dotted line) showing an increase of power in the broadband power (0.3-80 Hz). Middle: Average Vm traces during theta onset of an example depolarizing (red) cell, as well as the grand average over all events (grey) (n = 378, shaded areas represent ± sem). Bottom: Color plot showing the Vm of CA3 PCs triggered by transition to LIA normalized to 4 to 2 s before start of theta. Blue represents Vm hyperpolarization, red Vm depolarization. Each cell is represented in between black ticks. Cells are ordered depending on their average Vm modulation during theta, with depolarizing cells on top and hyperpolarizing cells at the bottom of the plot. Events within each cell are ordered by the time in which they occurred in the recording. (C) Relation between changes during theta and LIA in individual cells. Left: Change in membrane potential during theta vs during LIA. Right: Change in firing rate during theta vs during LIA. Each dot is one cell; edge color represents significant changes in Vm during theta and fill color represents significant changes in Vm during LIA (n = 32 cells).

**Figure S6.**
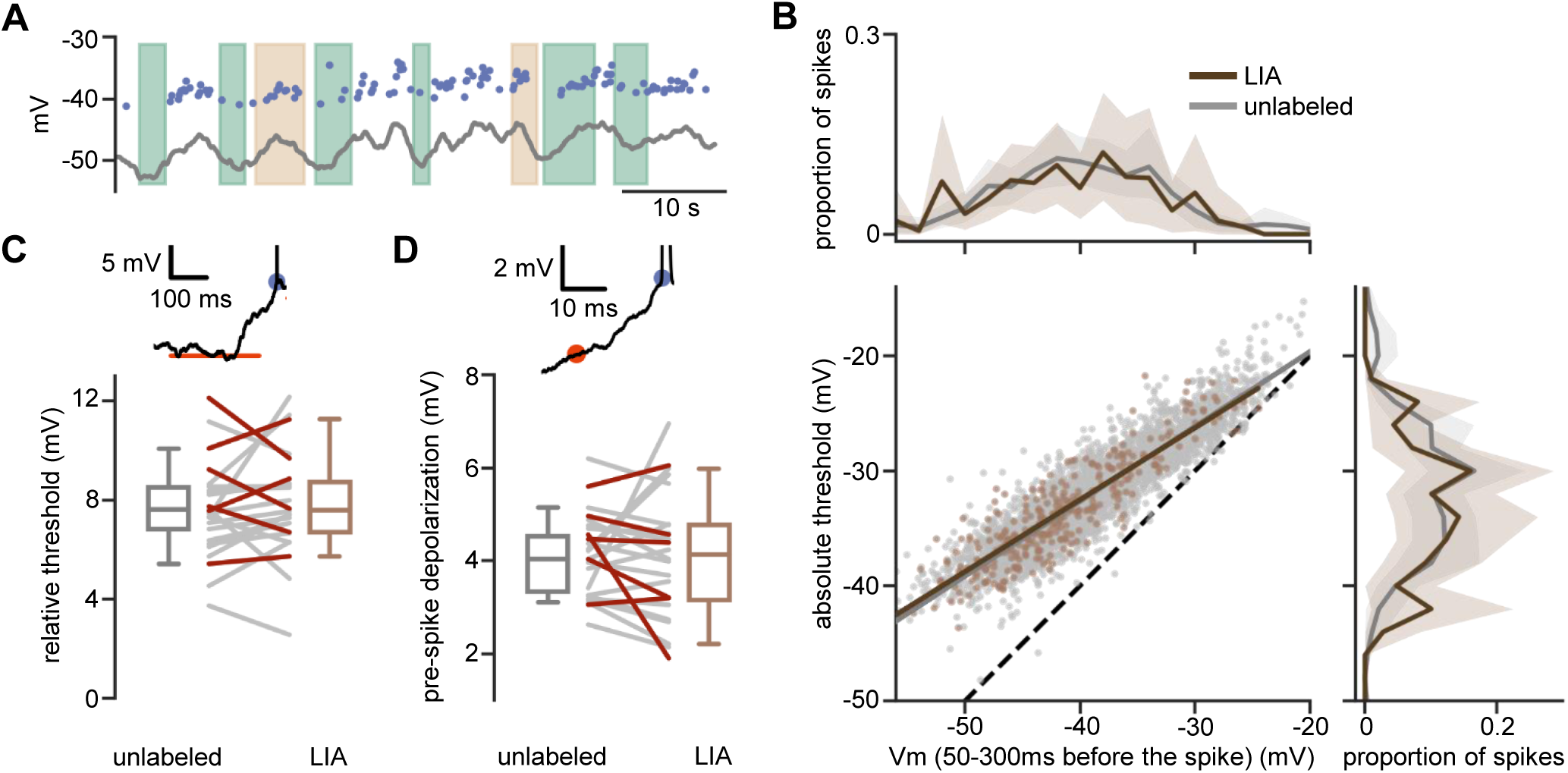
Dynamic changes in spike threshold relative to membrane potential during LIA. (A) Example trace showing smoothed Vm (grey line) and spike thresholds (blue dots). Shaded areas represent states. (B) Bottom left: Absolute spike threshold vs baseline Vm value 50-300 ms before the spike. Dots represent single spikes, color represents the state in which the spike occurs. The slopes and intercepts of the linear regressions were not significantly different between states (y = 0.65x – 6.6 for unlabeled [n = 3705 spikes], y = 0.63x – 7.5 for theta [n = 327 spikes], change in slope paired test over cells: p = 0.109, change in intercept paired test over cells: p = 0.092). Top: Histograms of the baseline Vm prior to spikes, separated by state and averaged over cells (-38.0 ± 4.9 mV [n = 31 cells] for unlabeled, -38.9 ± 4.4 mV [n = 25 cells] for LIA, paired test over cells: p = 0.005). Right: Histograms of the spike threshold, separated by states and averaged over cells (-30.9 ± 4.0 mV [n = 31 cells] for unlabeled, -32.7 ± 4.4 mV [n = 25 cells] for LIA, paired test over cells: p = 0.001). (C) Top: example spike (truncated) showing the absolute spike threshold (blue dot) and the Vm 500-300 ms before the spike (orange line). Bottom: Relative threshold (difference between absolute spike threshold and baseline Vm) for each cell during theta and unlabeled states. Colored lines represent cells that significantly depolarize (red) during LIA. There was no significant difference in the relative threshold over states (7.6 ± 1.6 mV [n = 24 cells] for LIA, 7.6 ± 1.3 mV [n = 31 cells] for unlabeled, p = 0.182). (D) Top: example spike (truncated) showing the absolute spike threshold (blue dot) and the Vm 20 ms before the spike (orange dot). Bottom: Same as (C) but for the pre-spike depolarization. There was no significant difference in the pre-spike depolarization over states (4.1 ± 1.1 mV [n = 24 cells] for LIA, 4.0 ± 0.6 mV [n = 31 cells] for unlabeled, p = 0.490).

**Figure S7.**
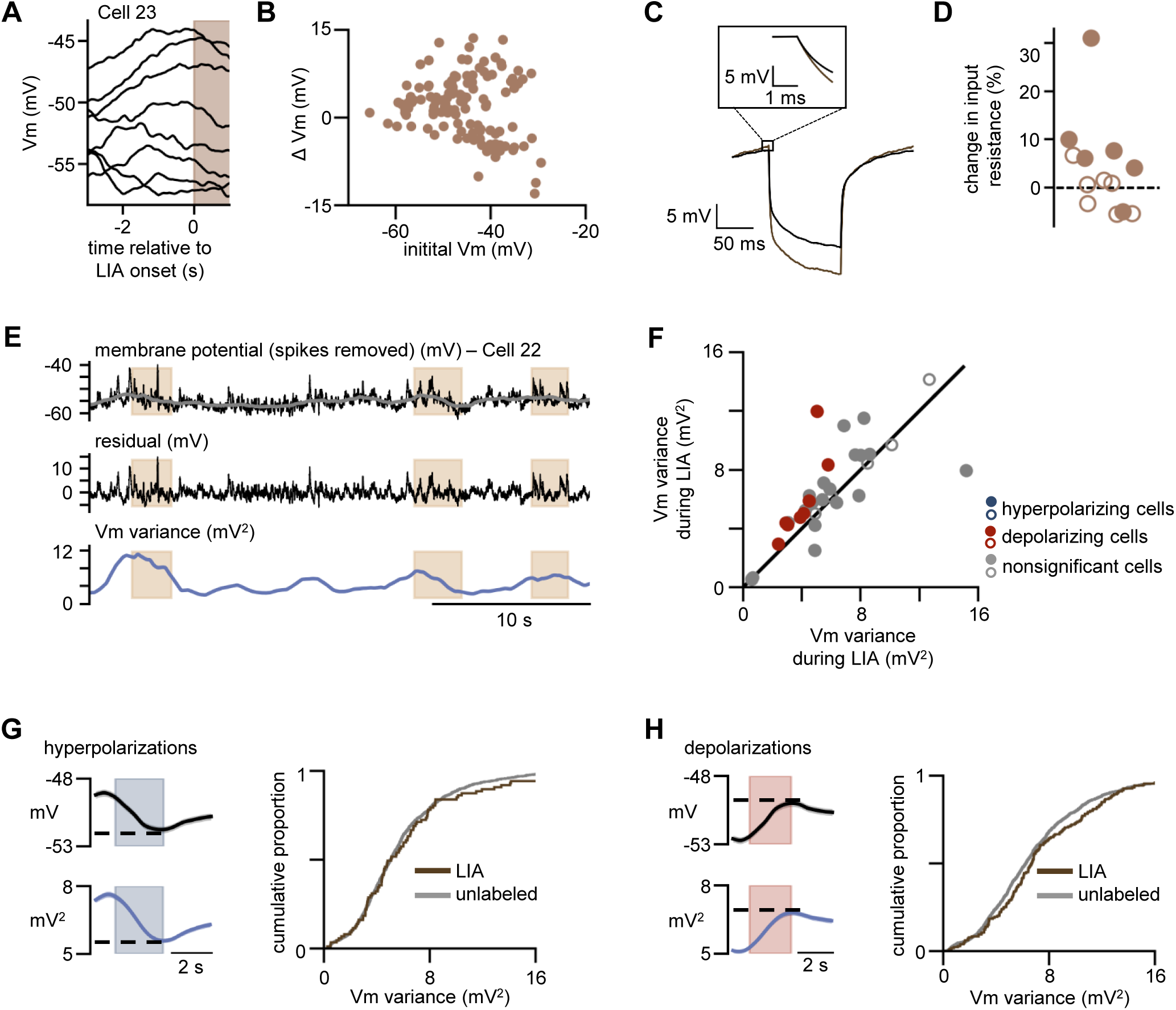
LIA-associated changes in membrane potential and firing rate occur with changes in input resistance and membrane potential variability. (A) Example traces showing Vm modulation during LIA for different initial Vm values in a single cell. (B) Vm changes during LIA (ΔVm) versus the initial membrane potential (initial Vm) of the cell [n = 122 events]. (C) Average voltage response to current pulses (-100 pA, 100 ms) in an example cell during LIA (brown) and unlabeled (black). Inset: zoom in at the onset of the current pulse. (D) Percentage change in input resistance during LIA compared to unlabeled (1.5% ± 6.4%, p = 0.874, [n = 13 cells]). Filled circles represent cells that had a significant difference in input resistance between LIA and unlabeled states [n = 6 cells]. (E) Processing of Vm trace to measure its variance. Top: downsampled Vm after spike removal of a CA3 PC with shading indicating LIA (brown). The grey superimposed trace is after smoothing with a window of 2 seconds. The smoothed Vm is subtracted from the spike-removed Vm to obtain the residual Vm (middle). The variance of the residual Vm trace is calculated over 1-second windows (bottom). (F) Scatter plot showing Vm variance of each cell during LIA versus nonLIA (n = 32 cells). Over the population of cells, there was a significant increase in variance during LIA (5.2 ± 2.2 mV^2^ for nonLIA, 6.1 ± 2.4 mV^2^ for LIA, p = 0.034). Color represents direction of modulation during LIA (blue: significantly hyperpolarizing cell: red: significantly depolarizing cell: grey: nonsignificant change), and filled dots represent a significant difference in variance between LIA and nonLIA. (G) Left: Average of all hyperpolarizing events Vm (top, black) and variance (bottom, blue, n = 1625 events). Dotted line corresponds to the value taken at the end of the change in Vm for each event. Right: Cumulative proportion of variance during LIA (brown) and unlabeled (grey) hyperpolarizing events. Vm variance is not significantly different between hyperpolarizations that occur during LIA and those that occur during unlabeled states (5.1 ± 3.2 mV^2^ [n = 88 events] for LIA, 5.1 ± 2.8 mV^2^ [n = 1177 events] for unlabeled, p = 0.953). (H) Left: Average of all depolarizing events Vm (top, black) and variance (bottom, blue, n = 1719 events). Dotted line corresponds to the value taken at the end of the change in Vm for each event. Right: Cumulative proportion of variance during theta and unlabeled depolarizing events. Vm variance is not different between depolarizations occurring during LIA and unlabeled states (6.6 ± 3.6 mV^2^ [n = 184 events] for LIA, 6.1 ± 3.5 mV^2^ [n = 1399 events] for unlabeled, p = 0.26).

**Figure S8.**
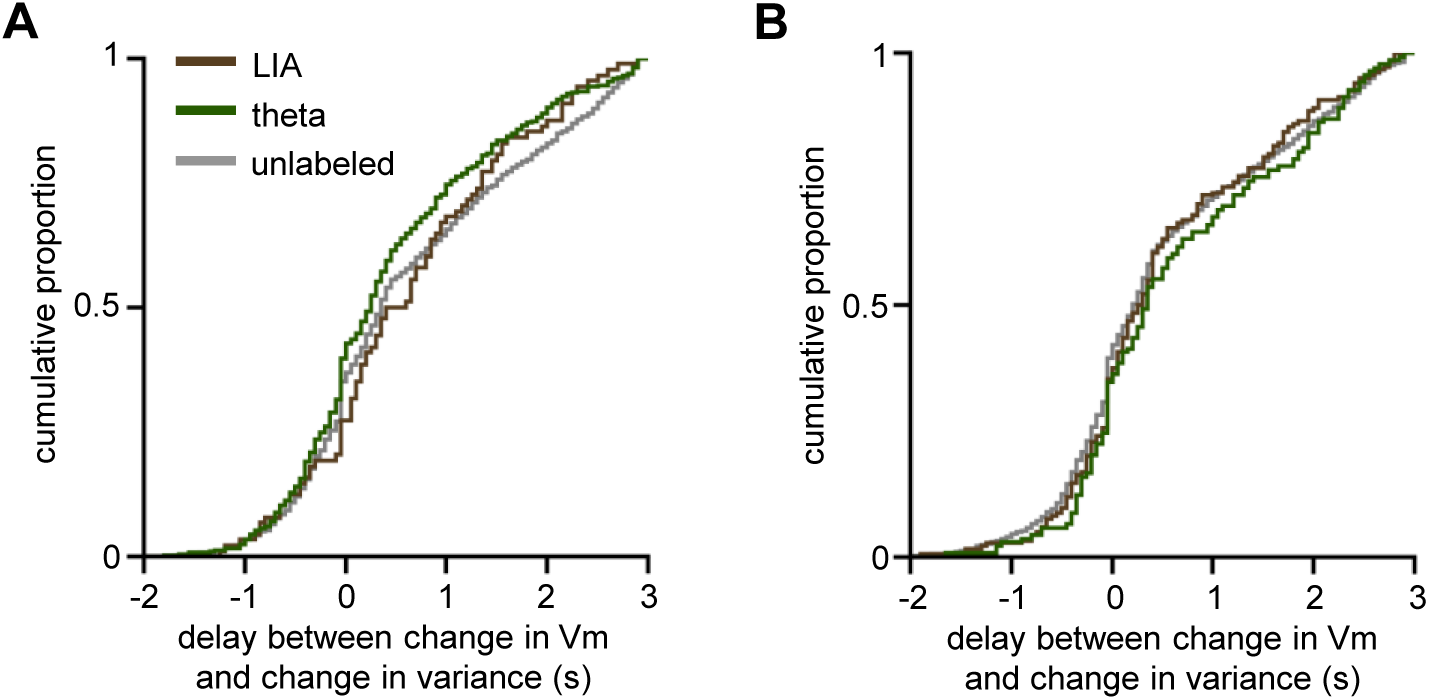
Nonzero delay between decrease in Vm and decrease in variance is not state-dependent. (A) Cumulative distribution of the lag between the start of the hyperpolarization and the start of the decrease in variance. Hyperpolarizations that coincide with the onset of theta (green), LIA (brown), and all other times (grey) are separated for comparison. All groups have a median significantly greater than zero (0.3 ± 0.8 s [n = 363 hyperpolarizations], p < 0.001 for theta, 0.6 ± 0.8 s [n = 88 hyperpolarizations], p = 0.001 for LIA, 0.4 ± 0.9 s [n = 1177 hyperpolarizations], p < 0.001 for unlabeled). Kruskall-Wallace suggests a significant difference (p = 0.030), but post-hoc bootstrapping does not reveal any significant differences (p = 0.067 for theta vs. unlabeled, p = 0.213 for LIA vs. unlabeled). (B) Same as in (A), but for depolarizations. All groups have a median significantly greater than zero (0.4 ± 0.9 s [n = 139 depolarizations], p < 0.001 for theta, 0.3 ± 0.8 s [n = 184 depolarizations], p < 0.001 for LIA, 0.3 ± 0.9 s [n = 1399 depolarizations], p < 0.001 for unlabeled). The delay was not different between states (Kruskal-Wallace, p = 0.236).

## References

Adam, Y., Kim, J.J., Lou, S., Zhao, Y., Xie, M.E., Brinks, D., Wu, H., Mostajo-Radji, M.A., Kheifets, S., Parot, V., et al. (2019). Voltage imaging and optogenetics reveal behaviour-dependent changes in hippocampal dynamics. Nature.

Atallah, B. V, and Scanziani, M. (2009). Instantaneous modulation of gamma oscillation frequency by balancing excitation with inhibition. Neuron 62, 566–577.

Azouz, R., and Gray, C.M. (2003). Adaptive coincidence detection and dynamic gain control in visual cortical neurons in vivo. Neuron 37, 513–523.

Bennett, C., Arroyo, S., and Hestrin, S. (2013). Subthreshold mechanisms underlying state-dependent modulation of visual responses. Neuron 80, 350–357.

Berry, S.D., and Thompson, R.F. (1978). Prediction of learning rate from the hippocampal electroencephalogram. Science 200, 1298–1300.

Bittner, K.C., Grienberger, C., Vaidya, S.P., Milstein, A.D., Macklin, J.J., Suh, J., Tonegawa, S., and Magee, J.C. (2015). Conjunctive input processing drives feature selectivity in hippocampal CA1 neurons. Nat. Neurosci. 18, 1133–1142.

Buzsáki, G. (1986). Hippocampal sharp waves: their origin and significance. Brain Res. 398, 242–252.

Buzsáki, G. (2002). Theta oscillations in the hippocampus. Neuron 33, 325–340.

Buzsáki, G. (2015). Hippocampal sharp wave-ripple: A cognitive biomarker for episodic memory and planning. Hippocampus 25, 1073–1188.

Buzsáki, G., and Moser, E.I. (2013). Memory, navigation and theta rhythm in the hippocampal-entorhinal system. Nat. Neurosci. 16, 130–138.

Carr, M.F., Jadhav, S.P., and Frank, L.M. (2011). Hippocampal replay in the awake state: A potential substrate for memory consolidation and retrieval. Nat. Neurosci. 14, 147–153.

Cohen, J.D., Bolstad, M., and Lee, A.K. (2017). Experience-dependent shaping of hippocampal CA1 intracellular activity in novel and familiar environments. Elife 6, 1–27.

Crochet, S., and Petersen, C.C.H. (2006). Correlating whisker behavior with membrane potential in barrel cortex of awake mice. Nat. Neurosci. 9, 608–610.

Eichenbaum, H. (2016). Still searching for the engram. Learn. Behav. 44, 209–222.

Ekstrom, A.D., Caplan, J.B., Ho, E., Shattuck, K., Fried, I., and Kahana, M.J. (2005). Human hippocampal theta activity during virtual navigation. Hippocampus 15, 881–889.

English, D.F., Peyrache, A., Stark, E., Roux, L., Vallentin, D., Long, M.A., Buzsaki, G., and Buzsáki, G. (2014). Excitation and Inhibition Compete to Control Spiking during Hippocampal Ripples: Intracellular Study in Behaving Mice. J Neurosci 34, 16509–16517.

Epsztein, J., Brecht, M., and Lee, A.K. (2011). Intracellular Determinants of Hippocampal CA1 Place and Silent Cell Activity in a Novel Environment. Neuron 70, 109–120.

Foster, D.J., and Wilson, M.A. (2006). Reverse replay of behavioural sequences in hippocampal place cells during the awake state. Nature 440, 680–683.

Fuhrmann, F., Justus, D., Sosulina, L., Kaneko, H., Beutel, T., Friedrichs, D., Schoch, S., Schwarz, M.K., Fuhrmann, M., and Remy, S. (2015). Locomotion, Theta Oscillations, and the Speed-Correlated Firing of Hippocampal Neurons Are Controlled by a Medial Septal Glutamatergic Circuit. Neuron 86, 1253–1264.

Gan, J., Weng, S., Pernía-Andrade, A.J., Csicsvari, J., and Jonas, P. (2017). Phase-Locked Inhibition, but Not Excitation, Underlies Hippocampal Ripple Oscillations in Awake Mice In Vivo. Neuron 93, 1–7.

Green, J.D., and Arduini, A.A. (1954). Hippocampal electrical activity in arousal. J. Neurophysiol. 17, 533–557.

Gupta, A.S., Van Der Meer, M.A.A., Touretzky, D.S., and Redish, A.D. (2012). Segmentation of spatial experience by hippocampal theta sequences. Nat. Neurosci. 15, 1032–1039.

Harvey, C.D., Collman, F., Dombeck, D.A., and Tank, D.W. (2009). Intracellular dynamics of hippocampal place cells during virtual navigation. Nature 461, 941–946.

Hulse, B.K., Moreaux, L.C., Lubenov, E. V, Siapas Correspondence, A.G., and Siapas, A.G. (2016). Membrane Potential Dynamics of CA1 Pyramidal Neurons during Hippocampal Ripples in Awake Mice. Neuron 89, 800–813.

Hulse, B.K., Lubenov, E. V., and Siapas, A.G. (2017). Brain State Dependence of Hippocampal Subthreshold Activity in Awake Mice. Cell Rep. 18, 136–147.

Jutras, M.J., Fries, P., and Buffalo, E.A. (2013). Oscillatory activity in the monkey hippocampus during visual exploration and memory formation. Proc. Natl. Acad. Sci. U. S. A. 110, 13144– 13149.

Kandel, E.R., Spencer, W.A., and Brinley, F.J. (1961). ELECTROPHYSIOLOGY OF HIPPOCAMPAL NEURONS: I. SEQUENTIAL INVASION AND SYNAPTIC ORGANIZATION. J. Neurophysiol. 24, 225–242.

Kesner, R.P., and Rolls, E.T. (2015). A computational theory of hippocampal function, and tests of the theory: New developments. Neurosci. Biobehav. Rev. 48, 92–147.

Komisaruk, B.R. (1970). Synchrony between limbic system theta activity and rhythmical behavior in rats. J. Comp. Physiol. Psychol. 70, 482–492.

Kowalski, J., Gan, J., Jonas, P., and Pernía-Andrade, A.J. (2016). Intrinsic membrane properties determine hippocampal differential firing pattern in vivo in anesthetized rats. Hippocampus 26, 668–682.

Landfield, P.W., Mcgaugh, J.L., and Tusa, R.J. (1972). Theta rhythm: A temporal correlate of memory storage processes in the rat. Science (80-.). 175, 87–89.

Lee, A.K., Epsztein, J., and Brecht, M. (2009). Head-anchored whole-cell recordings in freely moving rats. Nat. Protoc. 4, 385–392.

Leutgeb, S. (2004). Distinct Ensemble Codes in Hippocampal Areas CA3 and CA1. Science (80-.). 305, 1295–1298.

Leutgeb, J.K., Leutgeb, S., Moser, M.-B., and Moser, E.I. (2007). Pattern separation in the dentate gyrus and CA3 of the hippocampus. Science 315, 961–966.

Lin, P., Asinof, S.K., Edwards, N.J., and Isaacson, J.S. (2019). Arousal regulates frequency tuning in primary auditory cortex. Proc. Natl. Acad. Sci. 0, 201911383.

Macrides, F., Eichenbaum, H.B., and Forbes, W.B. (1982). Temporal relationship between sniffing and the limbic theta rhythm during odor discrimination reversal learning. J. Neurosci. 2, 1705–1717.

Maier, N., Tejero-Cantero, Á., Dorrn, A.L., Winterer, J., Beed, P.S., Morris, G., Kempter, R., Poulet, J.F.A., Leibold, C., and Schmitz, D. (2011). Coherent Phasic Excitation during Hippocampal Ripples. Neuron 72, 137–152.

Marder, E., O’Leary, T., and Shruti, S. (2014). Neuromodulation of Circuits with Variable Parameters: Single Neurons and Small Circuits Reveal Principles of State-Dependent and Robust Neuromodulation. Annu. Rev. Neurosci. 37, 329–346.

Margrie, T.W., Brecht, M., and Sakmann, B. (2002). In vivo, low-resistance, whole-cell recordings from neurons in the anaesthetized and awake mammalian brain. Pflugers Arch. Eur. J. Physiol. 444, 491–498.

Marr, D. (1971). Simple Memory: A Theory for Archicortex. Philos. Trans. R. Soc. B Biol. Sci. 262, 23–81.

McGinley, M.J., Vinck, M., Reimer, J., Batista-Brito, R., Zagha, E., Cadwell, C.R., Tolias, A.S., Cardin, J.A., and McCormick, D.A. (2015). Waking State: Rapid Variations Modulate Neural and Behavioral Responses. Neuron 87, 1143–1161.

Mizuseki, K., Sirota, A., Pastalkova, E., and Buzsáki, G. (2009). Theta Oscillations Provide Temporal Windows for Local Circuit Computation in the Entorhinal-Hippocampal Loop. Neuron 64, 267–280.

Nakazawa, K., Sun, L.D., Quirk, M.C., Rondi-Reig, L., Wilson, M.A., and Tonegawa, S. (2003). Hippocampal CA3 NMDA receptors are crucial for memory acquisition of one-time experience. Neuron 38, 305–315.

O’Keefe, J. (1976). Place units in the hippocampus of the freely moving rat. Exp. Neurol. 51, 78– 109.

Polack, P.O., Friedman, J., and Golshani, P. (2013). Cellular mechanisms of brain state-dependent gain modulation in visual cortex. Nat. Neurosci. 16, 1331–1339.

Poulet, J.F. a, and Petersen, C.C.H. (2008). Internal brain state regulates membrane potential synchrony in barrel cortex of behaving mice. Nature 454, 881–885.

Ramirez-Villegas, J.F., Logothetis, N.K., and Besserve, M. (2015). Diversity of sharp-wave– ripple LFP signatures reveals differentiated brain-wide dynamical events. Proc. Natl. Acad. Sci. 112, E6379–E6387.

Rebola, N., Carta, M., and Mulle, C. (2017). Operation and plasticity of hippocampal CA3 circuits: Implications for memory encoding. Nat. Rev. Neurosci. 18, 209–221.

Schneider, D.M., Nelson, A., and Mooney, R. (2014). A synaptic and circuit basis for corollary discharge in the auditory cortex. Nature 513, 189–194.

Taxidis, J., Anastassiou, C.A., Diba, K., and Koch, C. (2015). Local Field Potentials Encode Place Cell Ensemble Activation during Hippocampal Sharp Wave Ripples. Neuron 87, 590–604.

Vanderwolf, C.H. (1969). Hippocampal electrical activity and voluntary movement in the rat. Electroencephalogr. Clin. Neurophysiol. 26, 407–418.

Wixted, J.T., Squire, L.R., Jang, Y., Papesh, M.H., Goldinger, S.D., Kuhn, J.R., Smith, K.A., Treiman, D.M., and Steinmetz, P.N. (2014). Sparse and distributed coding of episodic memory in neurons of the human hippocampus. Proc. Natl. Acad. Sci. 111, 9621–9626.

Zucca, S., Griguoli, M., Malézieux, M., Grosjean, N., Carta, M., and Mulle, C. (2017). Control of Spike Transfer at Hippocampal Mossy Fiber Synapses In Vivo by GABA A and GABA B Receptor-Mediated Inhibition. J. Neurosci. 37, 587–598.

